# Studying the dawn of *de novo* gene emergence in mice reveals fast integration of new genes into functional networks

**DOI:** 10.1101/510214

**Authors:** Chen Xie, Cemalettin Bekpen, Sven Künzel, Maryam Keshavarz, Rebecca Krebs-Wheaton, Neva Skrabar, Kristian Ullrich, Diethard Tautz

## Abstract

The *de novo* emergence of new transcripts has been well documented through genomic analyses. However, a functional analysis, especially of very young protein-coding genes, is still largely lacking. Here we focus on three loci that have evolved from previously intergenic sequences in the house mouse *(Mus musculus)* and are not present in its closest relatives. We have obtained knockouts and analyzed their phenotypes, including a deep transcriptomic analysis, based on a dedicated power analysis. We show that the transcriptional networks are significantly disturbed in the knockouts and that all three genes have effects on phenotypes that are related to their expression patterns. This includes behavioral effects, skeletal differences and the regulation of the reproduction cycle in females. Substitution analysis suggests that all three genes have directly obtained an activity, without new adaptive substitutions. Our findings support the hypothesis that *de novo* genes can quickly adopt functions without extensive adaptation.

**Impact statement:** New protein-coding genes emerging out of non-coding sequences can become directly functional without signatures of adaptive protein changes

## Introduction

The evolution of new genes through duplication-divergence processes is well understood (Chen, Krinsky, & Long, 2013; Kaessmann, 2010; Long, Vankuren, Chen, & Vibranovski, 2013; Tautz & Domazet-Loso, 2011). But the evolution of new genes from non-coding DNA has long been only little considered (Tautz, 2014). However, with the increasing availability of comparative genome data from closely related species, more and more cases of unequivocal *de novo* transcript emergence have been described (McLysaght & Hurst, 2016; Schloetterer, 2015; Tautz, 2014; Tautz & Domazet-Loso, 2011). These analyses have shown that *de novo* transcript origination is a very active process in virtually all evolutionary lineages. A comparative analysis of closely related mouse species has even suggested that virtually the whole genome is “scanned” by transcript emergence and loss within about 10 million years of evolutionary history (Neme & Tautz, 2016).

But unlike the detection of the transcriptional and translational expression of *de novo* genes, functional studies of such genes have lacked behind. In yeast, the *de novo* evolved gene *BSC4* was found to be involved in DNA repair (Cai, Zhao, Jiang, & Wang, 2008) and *MDF1* (D. Li et al., 2010; D. Li, Yan, Lu, Jiang, & Wang, 2014) was found to suppress mating and promote fermentation. Knockdown of candidates of *de novo* genes in *Drosophila* have suggested effects on viability and fertility (Chen, Zhang, & Long, 2010; Reinhardt et al., 2013). However, in each of these cases, the genes were already relatively old, especially when taking the short generation times of these organisms into account. The most details for a very recent *de novo* evolved gene are so far available for *Pldi* in mice, which emerged 2.5-3.5 million years ago. In this case the knockout was shown to affect sperm motility and testis weight. But *Pldi* codes for a long non-coding RNA, not for a protein (Heinen, Staubach, Haming, & Tautz, 2009). Here, we focus on protein coding genes that have emerged less than 1.5 million years ago in the lineage towards the house mouse *(Mus musculus).*

There is abundant transcription of non-coding regions in vertebrate genomes (Consortium, 2012; Consortium et al., 2007; Neme & Tautz, 2016). Hence, the raw material for new genes is present at any time and most of these transcripts have at least short open reading frames (ORFs). Analysis of ribosome profiling data has shown that these are often translated (Ruiz-Orera, Messeguer, Subirana, & Alba, 2014; Ruiz-Orera, Verdaguer-Grau, Villanueva-Canas, Messeguer, & Alba, 2018), implying that many peptides derived from essentially random sequences can continuously be “tested” by evolution. If such a peptide conveys even a small evolutionary advantage, it is expected to come initially under stabilizing selection and eventually also under positive selection after acquiring further mutations. If it conveys a disadvantage, it should come under negative selection and should quickly be lost. In case it is evolutionary neutral, *i.e*., has no effect on the phenotype, it could still stay in the gene pool for some time, until a random disabling mutation occurs and becomes fixed in the population. Hence, for the youngest genes it is particularly important to show that they have effects on phenotypes, *i.e.,* they are not simply neutral bystanders.

Expression of random peptides in *E. coli* has shown that the majority is indeed not neutral, but conveys a growth disadvantage or advantage to the cells (Neme, Amador, Yildirim, McConnell, & Tautz, 2017). However, the conclusion of whether such peptides can convey indeed a direct advantage has been challenged (Knopp & Andersson, 2018; Tautz & Neme, 2018). Hence, it is of major interest to ask for very recently evolved protein-coding transcripts, whether these have already become integrated into regulatory networks and whether they have effects on phenotypes. It is important to study them at the “dawn” of gene emergence, *i.e.,* to capture them before further adaptation has taken place.

Using mouse as a model system for studying *de novo* gene evolution has the advantage that organ-specific, morphological and behavioral effects can be studied. The latter is of special relevance, since a large fraction of the *de novo* genes are initially expressed in the brain, possibly because they are somewhat shielded from the adaptive immune system (Bekpen, Xie, & Tautz, 2018). Further, a large diversity of recently differentiated populations and subspecies is available for mice, allowing to trace even very recent evolutionary events.

Here, we have generated a list of over one hundred candidate proteins that have evolved in the lineage of mice, after they split from rats. We show that most of these are translated, as inferred from ribosome profiling data, as well as mass spectrometry data. From this list, we have chosen three genes that have emerged particularly recent and subjected them to extensive molecular and phenotypic analysis. We conclude that all three of them have functions that would have been present from the time onwards at which they were born, without measurable further adaptation. These results support the notion that random peptide sequences have a good probability for conveying evolutionarily relevant functions.

## Results

### Recently evolved de novo genes in the mouse genome

To identify candidates for recently evolved *de novo* genes, we have applied a combined phylostratigraphy and synteny-based approach. Note that while the phylostratigraphy based approach was criticized to potentially include false positives (Moyers & Zhang, 2015), we have shown that the problem is relatively small and that it is in particularly not relevant for the most recently diverged lineages within which *de novo* gene evolution is traced (Domazet-Loso et al., 2017). We were able to identify 119 predicted protein-coding genes from intergenic regions that occur only in the mouse genome, but not in rats or humans (Figure 1 - figure supplement 1). We re-assembled their transcriptional structures and estimated their expression levels using available ENCODE RNA-Seq data in 35 tissues (Figure 1). To validate that their predicted ORFs are indeed translated, we have searched ribosome profiling and peptide mass spectrometry datasets (Figure 1 - figure supplement 1). We found for 110 out of the 119 candidate genes direct evidence for translation.

Expression of these genes is found throughout all tissues analyzed, with notable differences. Testis and brain express the highest fraction, while the digestive system and liver express the lowest fraction (Figure 1A). Expression levels of these genes are generally lower than those of other genes (FPKM medians: 0.63 vs. 8.18; two-tailed Wilcoxon rank sum test, P-Value < 2.2 × 10^−16^; Figure 1C). Most overall molecular patterns are similar to previous findings (Neme & Tautz, 2013; Schmitz, Ullrich, & Bornberg-Bauer, 2018). They have fewer exons (medians: 2 vs. 7; two-tailed Wilcoxon rank sum test, P-Value < 2.2 × 10” ^16^) and fewer coding exons than other genes (medians: 1 vs. 6; two-tailed Wilcoxon rank sum test, P-Value < 2.2 × 10^−16^). The lengths of their proteins are shorter than those of other proteins (medians: 125 vs. 397; two tailed Wilcoxon rank sum test, P-Value < 2.2 × 10^−16^). However, their proteins are predicted to be less disordered than other proteins (medians: 0.20 vs. 0.27; two-tailed Wilcoxon rank sum test, P-Value = 0.0024; Figure 1D) and equally hydrophobic to other proteins (medians: 0.56 vs. 0.57; two-tailed Wilcoxon rank sum test, P-Value = 0.52; Figure 1E). Note that the two sets of values show a broad distribution.

### Genes for functional analyses

We selected three genes from the above list for in-depth analyses, including knockouts, transcriptomic studies and phenotyping (Table 1). For convenience we will call these genes in the following *“Unnamed de novo genes – Udng”, i.e., Udng1, Udng2,* and *Udng3,* but note that we propose new formal names in the discussion. The criteria for selecting these three genes were as follows: (i) they have clear transcriptional expression evidence, (ii) have at least two exons, (iii) their translation is supported by ribosome profiling and/or proteomic evidence and (iv) they are specific to the *M. musculus* lineage, *i.e.*, have emerged less than 1.5 million years ago (see below). Further, they cover also a range from low to high intrinsic structural disorder scores and hydrophobicities, as well as lower to higher expression levels (Figures 1C-E; Table 1).

**Figure 1.**
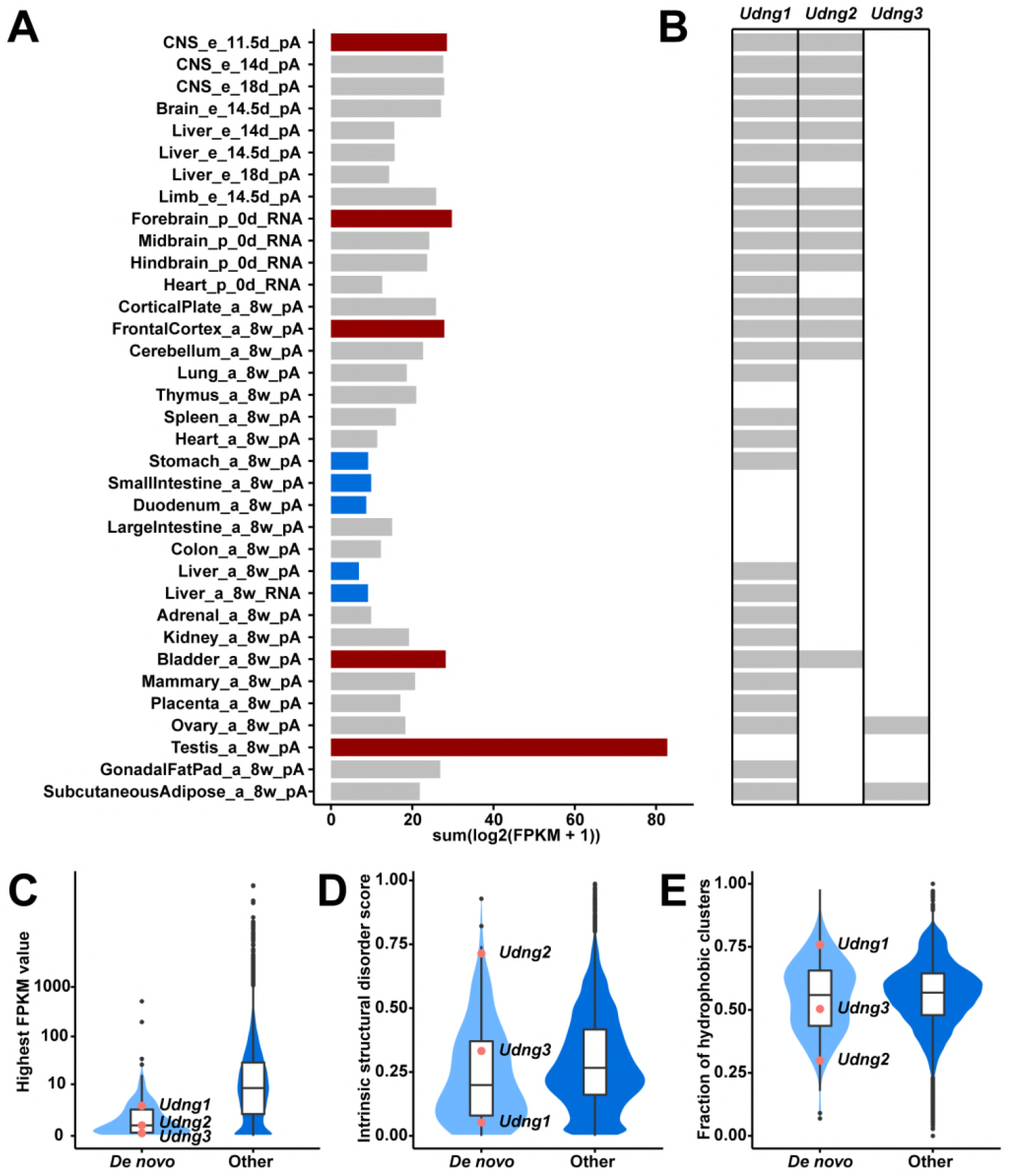
Transcriptional abundance and structural features of the 119 candidate *de novo* genes. (A) Transcriptional abundance in each tissue, represented as the sum of log transformed FPKM value of each transcript. Details on tissue designations and RNA samples are provided in Figure 1 - figure supplement 1. The five tissues with the highest fractions are highlighted in red and the lowest ones in blue. (B) Transcriptional abundance of the three genes studied here, *Udng1, Udng2,* and *Udng3* in each tissue. FPKM values greater than or equal to 0.1 are marked as gray, lower levels or absence in white. (C) Comparison of overall expression levels (represented as the highest FPKM values in the 35 tissues) between *de novo* and other protein-coding genes. (D) Comparison of averages of intrinsic structural disorder scores between *de novo* and other protein-coding genes. (E) Comparison of fractions of sequence covered by hydrophobic clusters between *de novo* and other protein-coding genes. The corresponding values for the three genes studied here (see Table 1) are indicated in the three violin plots.

**Table 1.**
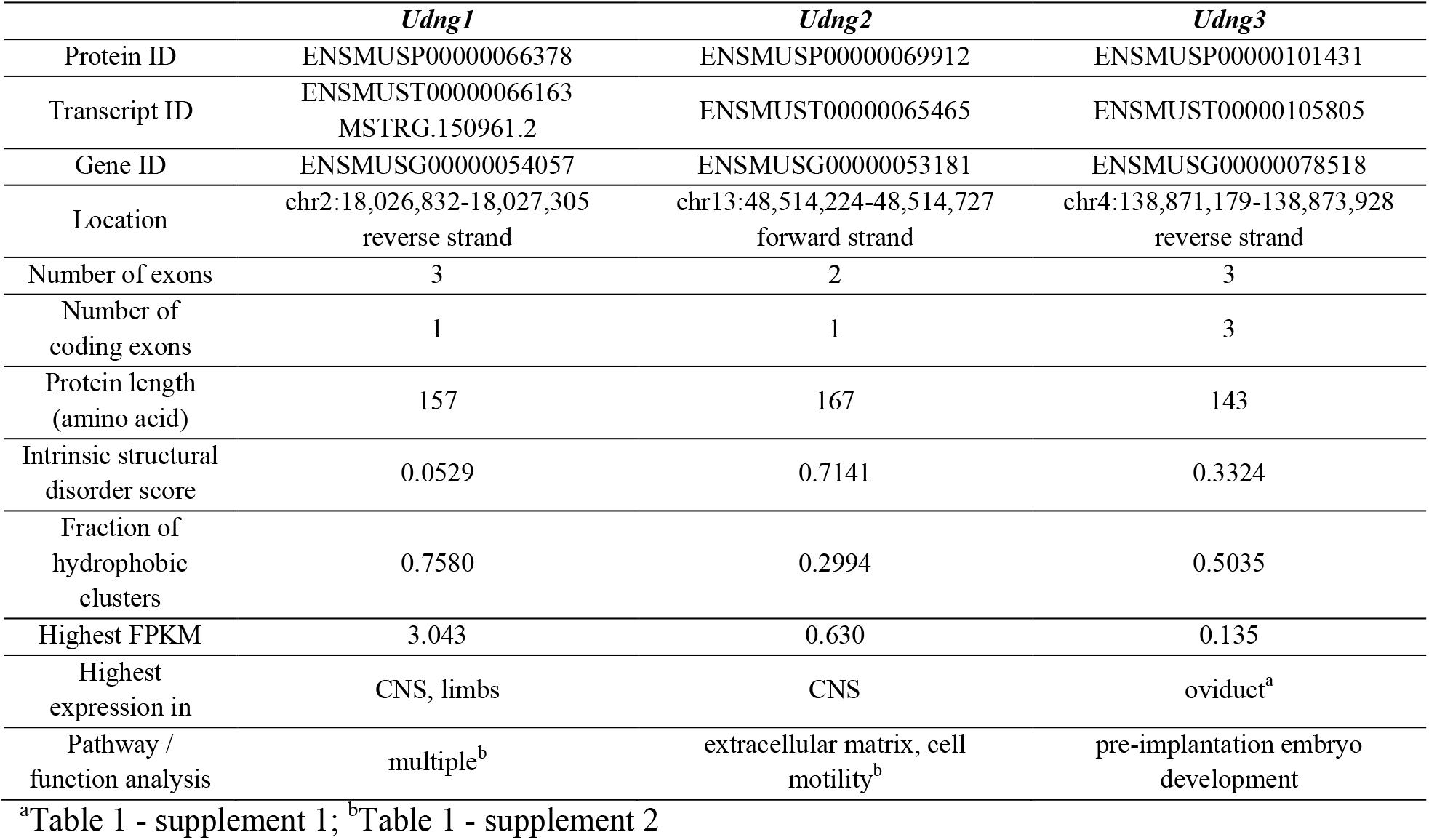
General information on the three genes selected for functional analyses.

*Udng1* shows a relatively high expression (up to FPKM 3) in multiple tissues, with the highest in brain tissues at different stages as well as in embryonic limbs (Figure 1B; Figure 1 - figure supplement 1).

*Udng2* shows on average a lower expression (up to FPKM 0.6), also mostly in brain tissues at different stages (Figure 1B; Figure 1 - figure supplement 1). *Udng3* is only expressed in two tissues, the ovary of 8 weeks old females (FPKM 0.135), as well as the subcutaneous adipose tissue of 8 weeks old animals (FPKM 0.115) (Figure 1B). Given that the ovary is a very small organ, with closely attached tissues, such as oviduct and gonadal fat pad, there could be contamination between these different tissue types. Hence, we were interested whether there is specificity for one of them. We used RT-PCR for the respective carefully prepared tissue samples for *Udng3* and a control gene *(Uba1)* and found that *Udng3* is not expressed in the ovary, but predominantly in the oviduct with only a weak signal from the adjacent fat pad (Table 1 - supplement 1).

### Evolutionary emergence of the three candidate genes

We used whole genome sequencing data (Harr et al., 2016) and Sanger sequencing data of PCR fragments from mouse populations, subspecies and related species to trace the emergence of the ORFs for the three candidate genes. We found the respective genomic regions covering the ORFs in all species analyzed, which include the wood mouse *Apodemus* that has split from the *Mus* lineage about 10 million years ago. However, in these more distant species, the reading frames are interrupted by early stop codons and/or non-frame indels. Full reading frames were only found in populations and subspecies of *M. musculus,* but not in *M. spretus* or *M. spicilegus* as the closest outgroups (Figure 2, Figure 2 – figure supplement 1). This implies that they have arisen after the split between these species and the *M. musculus* subspecies about 1.5 million years ago (Dejager, Libert, & Montagutelli, 2009). The *M. musculus* subspecies have split further into three major lineages, *M. m. castaneus, M. m. musculus* and *M. m. domesticus* about 0.5 million years ago (Figure 2). The three genes occur in at least two of these lineages (see below), *i.e.*, they are between 0.5 – 1.5 million years old.

**Figure 2.**
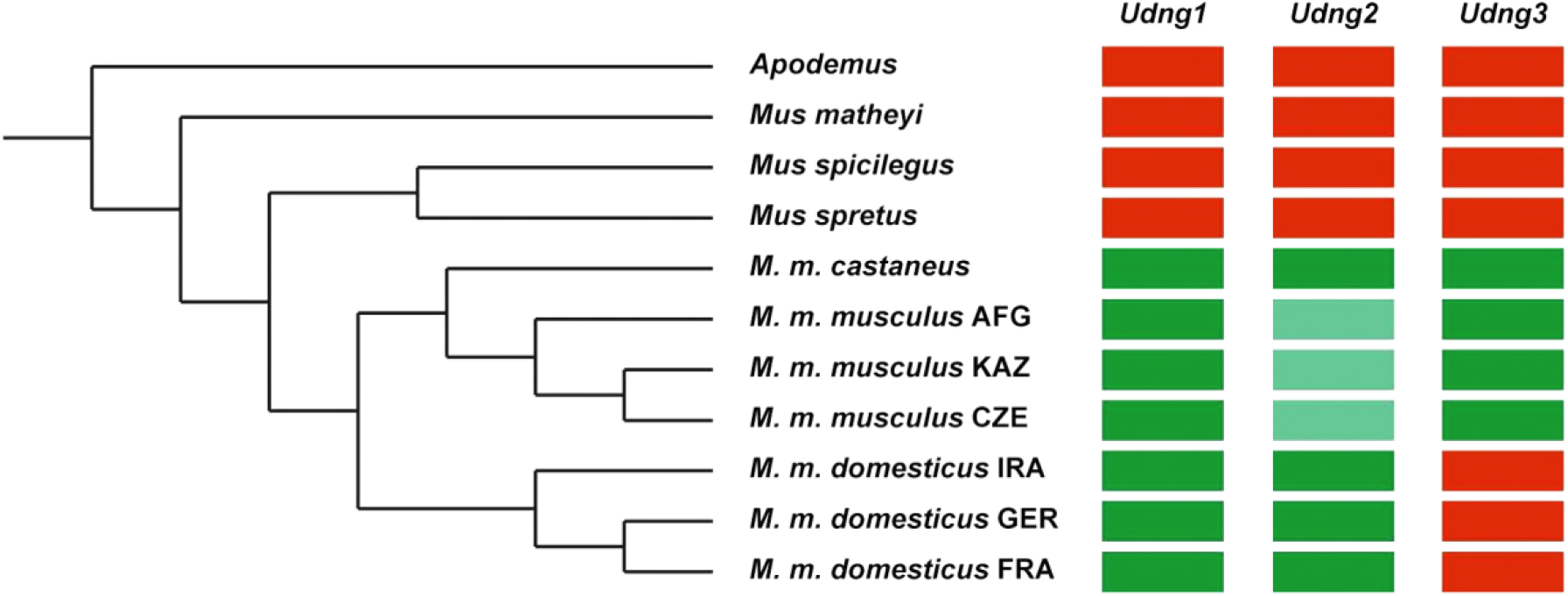
Emergence of the ORFs for the three genes. Left is the phylogenetic tree of the mouse species, subspecies *(Mus musculus* = *M. m.*), and the outgroup *Apodemus,* derived from whole genome sequence analyses (see Methods). Three populations each represent *M. m. musculus* (AFG = from Afghanistan, KAZ = from Kazakhstan, CZE = from Czech Republic) and *M. m. domesticus* (IRA = from Iran, FRA = from France, GER = from Germany). The right panel shows whether the ORF of each gene is intact or not. Red: not intact, green: intact, light green: almost intact, *i.e.*, secondary acquisition of a premature stop codon. The alignments of the coding sequences are provided in Figure 2 – figure supplement 1. The distance matrices are provided in Figure 2 – figure supplement 2.

*Udng1* occurs in all three subspecies and all analyzed populations. The same pattern is seen for *Udng2,* with the exception that the three *M. m. musculus* populations show a slightly shorter version (153 instead of 167 amino acids), due to a newly acquired premature stop codon (Figure 2 – figure supplement 1). *Udng3* is present in *M. m. castaneus* and *M. m. musculus,* while all three *M. m. domesticus* populations share a derived indel that disrupts its reading frame after 15 amino acids (Figure 2 – figure supplement 1).

None of the three gene regions show significant signatures of selection (TajD or F_ST_ analysis) in the population analyses provided in (Harr et al., 2016). Further, they show too few substitutions (Figure 2 - supplement 2) to allow a meaningful calculation of dN/dS ratios because of lack of power. To assess whether they show signs of an accelerated evolution after the acquisition of their ORFs, we have calculated the distances *(i.e.*, number of substitutions) within the tree of species analyzed. Using *M. matheyi* as the out-group, we can compare the average distances to the two species that show no ORF and should therefore evolve with an approximately neutral rate (*M. spretus* and *M. spicilegus* = non-coding group) with the average distances to the taxa that have the respective ORF *(M. m. castaneus, M. m. musculus* and *M. m. domesticus* = coding group) (see Figure 2 for these relationships). The latter should show on average more substitutions, if evolution was accelerated due to positive selection after the acquisition of the ORF. However, we find that this is not the case, the observed number of substitutions is very similar between both groups (Table 2). However, we noted that *Udng3* shows more substitutions for both groups. To obtain an estimate for the expected number of substitutions, we have used the average distances between the taxa derived from whole genome comparisons. These should reflect approximately the neutral rates, given that most of the genome is not expected to be subject to evolutionary constraints. The results are also provided in Table 2 (the full matrix of pairwise differences is included in Figure 2 – figure supplement 2). We find that *Udng1* and *Udng2* evolve at the expected average rate while *Udng3* is indeed faster than expected. Still, when testing observed versus expected values between each group for each locus, we find that none of them is significant (Table 2). Hence, in spite of the region specific rate differences, there are no signs that accelerated evolution through positive selection would have taken place after the acquisition of the ORFs in any of the three loci. However, we can not exclude that a selective sweep could have occurred at the time where the ORFs emerged, but this can not be traced anymore in todays populations.

**Table 2.**
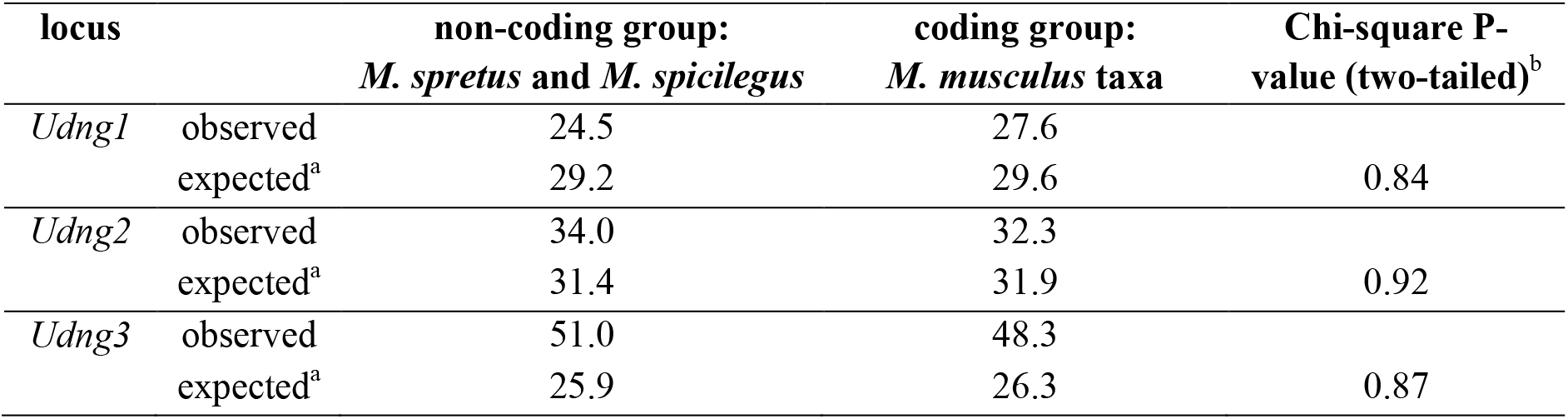
Average numbers of substitutions for each locus compared to *M. matheyi.*

### Generation of gene knockouts and power analysis

For the further functional characterization of the three genes, we obtained knockout lines. *Udng1* and *Udng2* represent constructs in which all or most of the ORFs were substituted by *lacZ*, *Udng3* was generated by creating a frame shift in the ORF through CRISPR/Cas9 mutagenesis. All three lines were homozygous viable and showed only subtle phenotypes (further details below). We were therefore interested in studying their impact on the transcriptional network in the tissues in which they are predominantly expressed. Given the recent evolution of the genes, one would expect only a small influence. Hence, we first did a power analysis to get an estimate on how deeply we can trace changes in the networks.

Several conditions have to be considered for such a power analysis. When using RNA-Seq read count (fragment count for paired-end sequencing) data, we assume (1) read counts follow a negative binomial distribution; (2) all samples are sequenced at the same depth; (3) significance level after Bonferroni adjusted is 0.05 and in total 15,000 genes are tested, *i.e.,* the significance level before adjustment is 3.3 × 10^−6^. The power to detect a differentially expressed gene can then be estimated by the given (1) sample size, (2) fold change between knockouts and wildtypes, (3) average read count, and (4) dispersion, which is the measurement of biological and technical variance considering the effect of mean read count (Figure 3A). Based on this a priori analysis, we used at least 10 biological replicates of knockouts and wildtypes, performed deep sequencing and minimized variance by using standardized rearing conditions for the mice, as well as standardized and parallel preparation and sequencing procedures. Under these conditions, it is expected to be possible to detect significant differences even when the fold-changes are as low as 1.05 to 1.25. We found that these expectations fitted well with our real data described below (Figure 3B and C, and Figure 3 – figure supplement 1).

**Figure 3.**
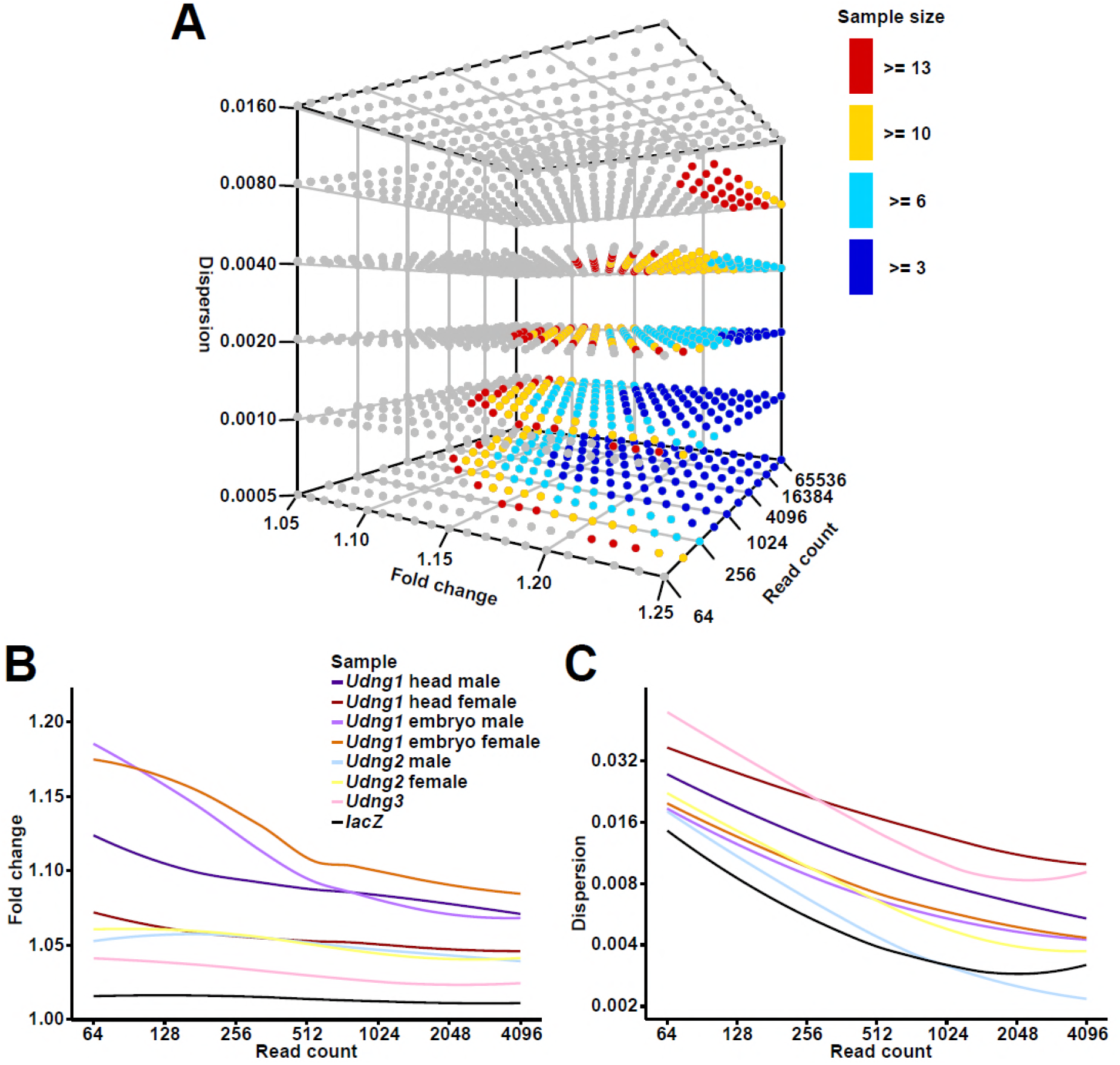
Power analysis and the comparison of the actual RNA-Seq datasets. (A) The theoretically estimated power for each combination of sample size, fold change, read count, and dispersion. The three axes represent fold change, read count, and dispersion separately. The grey dots represent power lower than 0.8, and the colored dots represent power greater than or equal to 0.8 under different sample sizes. (B and C) Curves of fold change (B) and dispersion (C) against read count from the actual RNA-Seq datasets, fitted with locally estimated scatterplot smoothing (LOESS) method. Values are taken from DESeq2 (read count as baseMean, fold change as 2^|log2FoldChange|^, and dispersion). Numeric details for the actual sample analysis are provided in Figure 3 – figure supplement 1.

### Controls

To assess whether any possible effects on the transcriptome could be caused by the expression of *lacZ* in *Udng1* and *Udng2,* we conducted a control experiment in cell culture. We transformed primary mouse embryonic fibroblasts with vectors expressing transcripts containing the *lacZ* ORF in forward and reverse direction. This was done in 10 parallels for each direction and RNA-Seq data were obtained for each of them after 48 hrs incubation *(i.e.,* transient expression). The expression of the transcripts including the *lacZ* ORF in the forward and the reverse directions were confirmed by the unique mapped reads. On average we could map 54.2 million unique reads per sample (range from 44.2 to 65.8 million reads). We did not detect any significantly differentially expressed genes in this experiment. This suggests that LacZ protein expression by itself does not result in traceable changes of the transcriptome. This conclusion applies of course only to this particular experiment and it could be useful to eventually repeat this in a whole mouse background. However, another control already inherent in our data is that in the RNA-Seq data of the heads of postnatal 0.5-day *Udng1* and *Udng2* male pups (see below). Both of these express *lacZ* but the sets of differentially expressed genes are different (they overlap only in 63 genes, whereby 79 would have been expected by chance).

The CRISPR/Cas9 experiment to generate our *Udng3* knockout line might have generated potential off-target mutations. In order to rule out this possibility, we performed whole genome sequencing on both animals of our founding pair. The female and male of our founding pair were selected from the first generation offspring of the mating among mosaic and wildtype mice which were directly developed from the zygotes injected. Each of them contained a 7-bp deletion allele and a wildtype allele. If there were any off-target sites, they should exist as heterozygous or homozygous indels or single nucleotide variants. However, in our genome sequencing results, we found no variant located in the 100 bp regions around the genome-wide 343 predicted off-target sites. Further, we manually checked the reads mapped to the regions around the top 20 predicted sites in both samples and none of them yielded an indication of variants.

In the light of these controls, we conclude that the effects shown for the knockouts in the following can indeed be ascribed to the knockouts themselves, rather than a confounding factor. We describe the results for each gene in turn.

### Udng1 knockout effect on the transcriptome

For *Udng1* the replacement construct removes the whole ORF. *Udng1* is broadly expressed across developmental stages and tissues (Figure 1B, Figure 1 - figure supplement 1). High expression in brain tissues is seen in embryos and pups and the limbs in embryos (Figure 3 – figure supplement 1). Hence, we used the heads of postnatal 0.5-day pups and 12.5-day whole embryos for RNA-Seq analysis.

We sequenced the heads of 10 postnatal 0.5-day pups from each of the four sex (female or male) and genotype (homozygous knockout or wildtype) combinations. On average, we could map 74.6 million unique reads for each sample (range from 59.3 to 89.4 million reads; Figure 3 – figure supplement 1).

First we examined whether the *Udng1* transcript was indeed lacking in the knockouts. This is the case: knockouts show no transcription, but wildtypes show clear transcription (Figure 3 – figure supplement 1). We also confirmed their genotypes by checking the level of *lacZ* expression (Figure 3 – figure supplement 1). We found 1,719 differentially expressed genes between male knockout and wildtype samples (DESeq2, adjusted P-Value ≤ 0.01, fold changes range from 0.649 to 1.36; Figure 3 – figure supplement 2). Interestingly, we found only one differentially expressed gene between females, *Udng1* itself (DESeq2, adjusted P-Value ≤ 0.01). This can be ascribed to a higher dispersion in the female samples (Figure 3C), which results in a loss of power. The reason for the higher dispersion in females in these samples is currently unclear. Functional enrichment analysis of the 1,718 differentially expressed genes (except for *Udng1* itself) in males revealed 501 distinct Gene Ontology functional terms and 137 distinct pathways (KOBAS, corrected P-Value ≤ 0.05; Table 1 - supplement 2).

RNA was also obtained from 10 to 14 12.5-day embryos of the four sex (female or male) and genotype (homozygous knockout or wildtype) combinations. On average, we could map 67.1 million unique reads per sample (range from 36.9 to 92.7 million reads; Figure 3 – figure supplement 1). Again we confirmed that the *Udng1* transcript was indeed lacking in the knockouts, and checked the level of *lacZ* expression (Figure 3 – figure supplement 1). We found 3,855 differentially expressed genes between male knockout and wildtype samples (DESeq2, adjusted P-Value ≤ 0.01, fold changes range from 0.533 to 1.59; Figure 3 – figure supplement 2) and 6,165 between females (DESeq2, adjusted P-Value ≤ 0.01, fold changes range from 0.531 to 1.56; Figure 3 – figure supplement 2). Among them, there are 2,998 shared between female and male samples. Functional enrichment analysis of the common differentially expressed genes revealed 583 distinct Gene Ontology functional terms and 137 distinct pathways (KOBAS, corrected P-Value ≤ 0.05; Table 1 - supplement 1). Among the 1,719 differentially expressed genes between male head samples and the 3,855 ones between male embryo samples, 418 are overlapping. In addition, there are 176 overlapping Gene Ontology functional terms and 17 overlapping pathways between the two datasets.

### Udng1 knockout effect on mouse behavior and limb length

The relatively high expression of *Udng1* in the CNS and the RNA-Seq results of the heads of postnatal pups indicate that it may have an effect on the behavior of the mice. We performed three standardized behavioral tests: elevated plus maze, open field, and novel object to test this possibility. We found a significant difference for the open field test with respect to total distance moved (nested ranks test, P-Value = 0.0023; Table 3; full data in Table 3 - supplement 1).

**Table 3.** Phenotyping results for *Udng1* and *Udng2.*

supplement 1 provides the details of the phenotype scores.
|  |  | *Udng1* |  |  |  | *Udng2* |  |  |  |
| --- | --- | --- | --- | --- | --- | --- | --- | --- | --- |
| Test | Parameter | N<sup>a</sup> | KO<sup>b</sup> | WT<sup>b</sup> | P-Value<sup>c</sup> | N<sup>a</sup> | KO<sup>b</sup> | WT<sup>b</sup> | P-Value<sup>c</sup> |
| Elevated plus maze | center time (%) | 40 | 11.9 | 10.8 | 0.19 | 36 | 10.8 | 14.8 | 0.029 |
|  | dark time (%) | 40 | 54.1 | 56.7 | 0.20 | 36 | 63.2 | 58.3 | 0.072 |
|  | light time (%) | 40 | 31.0 | 28.5 | 0.15 | 36 | 21.7 | 20.5 | 0.45 |
| Open field | wall time (%) | 40 | 51.4 | 44.7 | 0.24 | 12 | 58.6 | 49.5 | 0.29 |
|  | total distance (m) | 40 | 42.1 | 48.0 | 0.0023 | 12 | 31.7 | 35.0 | 0.29 |
| Novel object | first contact time (s) | 40 | 2.5 | 5.0 | 0.26 | 12 | 0.0 | 0.0 | 0.30 |
|  | object visits (N) | 40 | 4.0 | 3.0 | 0.14 | 12 | 0.0 | 0.0 | 0.39 |
|  | total distance (m) | 40 | 28.2 | 30.1 | 0.35 | 12 | 25.7 | 25.1 | 0.53 |
| Limb elements (length in mm) | humerus | 40 | 11.96 | 11.96 | 0.93 | n.d. |  |  |  |
|  | ulna | 40 | 13.86 | 13.83 | 0.37 | n.d. |  |  |  |
|  | metacarpal | 40 | 3.20 | 3.22 | 0.043 | n.d. |  |  |  |
|  | femur | 40 | 15.34 | 15.44 | 0.21 | n.d. |  |  |  |
|  | tibia | 40 | 17.37 | 17.21 | 0.072 | n.d. |  |  |  |
|  | metatarsal | 40 | 7.43 | 7.29 | 0.020 | n.d. |  |  |  |

<sup>a</sup>N = total number of individuals used, equally divided between knockouts and wildtypes.
<sup>b</sup>Medians across all individuals.
<sup>c</sup>P-Values for the behavior phenotypes were calculated using nested ranks tests representing a non-parametric linear mixed model; for the data having only one group, it is essentially identical to a one-tailed Wilcoxon rank sum test. For the limb length measurements we use a two-tailed Wilcoxon rank sum test.

Given that *Udng1* is also expressed in limbs, we asked whether there would also be differences in limb morphology. We scanned the skeletons of the respective wildtype and knockout mice and analyzed their bone lengths, following the procedures described in (Skrabar, Turner, Pallares, Harr, & Tautz, 2018). We found that the knockout mice had significantly longer metatarsals (two-tailed Wilcoxon rank sum test, P-Value = 0.020) and significantly shorter metacarpals (two-tailed Wilcoxon rank sum test, P-Value = 0.043), and in tendency also longer tibias (Table 3; full data in Table 3 - supplement 1)

This raises the question whether the limb length phenotype could cause the “distance moved” phenotype in the open field test (see above). However, given that “distance moved” was also recorded in the novel object test and showed no significant difference between WT and KO (see also discussion), we do not consider the small differences in limb length elements as factors that would impair movement. Hence, it is more likely that these phenotypes are independent of each other and relate to the different expression aspects in limbs and brains.

For *Udng2* the replacement construct removes 502 out of 504 base pairs of its ORF. *Udng2* is expressed in brain tissues at different stages (Figure 1B, Figure 1 - figure supplement 1) and we targeted the RNA-Seq analysis to the heads of postnatal 0.5-day pups. We sequenced the heads of 10 individuals each of the four sex (female or male) and genotype combinations (homozygous knockout or wildtype). On average, we could map 64.7 million unique reads for each sample (range from 57.0 to 74.4 million reads; Figure 3 – figure supplement 1). We confirmed that the *Udng2* transcript was indeed lacking in the knockouts, and checked the level of *lacZ* expression (Figure 3 – figure supplement 1). We found 1,399 differentially expressed genes between male knockout and wildtype samples (DESeq2, adjusted P-Value ≤ 0.01; fold changes range from 0.720 to 1.38; Figure 3 – figure supplement 2), but only 160 between females (DESeq2, adjusted P-Value ≤ 0.01; fold changes range from 0.757 to 1.33; Figure 3 – figure supplement 2). Similarly as seen in the *Udng1* analysis, we find a higher dispersion among the female samples that lowers the power of detection. Functional enrichment analysis of the differentially expressed genes in males reveals 306 distinct Gene Ontology functional terms and 14 pathways. All the pathways are related to extracellular matrix or cell motility functions (KOBAS, corrected P-Value ≤ 0.05; Table 1 - supplement 1).

### Udng2 knockout effect on mouse behavior

The RNA-Seq results of the heads of postnatal pups indicate that *Udng2* may be involved in mouse behavior too. We performed the same four behavioral tests as for *Udng1*. We found significant effects in the elevated plus maze test (Table 3 and Table 3 - supplement 1), but note that only fewer animals were available for the other tests. We found that knockout males stayed shorter in the center (nested ranks test, P-Value = 0.029), indicating a decision-making related phenotype (Cruz, Frei, & Graeff, 1994; Fernandes & File, 1996; Rodgers & Johnson, 1995) and they stayed longer in the dark arms (nested ranks test, P-Value = 0.072), indicating an anxiety related phenotype (Walf & Frye, 2007) (Table 3).

The *Udng3* knockout line was generated using CRISPR/Cas9 mutagenesis in a laboratory strain that is nominally derived from *M. m. domesticus* (C57BL/6N). As pointed out above, *M. m. domesticus* populations have already a disabling mutation for *Udng3.* However, C57BL/6N is known to carry also alleles from *M. m. musculus* (Yang et al., 2011) and the *Udng3* allele represents indeed the non-interrupted version that is found in *M. m. musculus* and *M. m. castaneus.* The CRISPR/Cas9 treatment introduced a 7-bp deletion at the beginning of the ORF (position 41-47) causing a frame shift and a premature stop codon in exon 2. Given the observation that *Udng3* is specifically expressed in adult oviducts (see above), we focused the RNA-Seq analysis on the oviducts of 12 knockout and 12 wildtype females (10-11 weeks old). There were on average 75.9 million unique mapped reads per sample (range from 57.5 to 93.0 million reads; Figure 3 – figure supplement 1). The genotypes of the 24 samples were further confirmed by the reads covering the sites in which the 7-bps deletion locates. In the initial analysis involving all samples, we found no differentially expressed gene between knockouts and wildtypes.

However, given that the expression in oviducts should be fluctuating according to estrous cycle, we clustered the transcriptomes of the individuals based on both principle component analysis (PCA) and hierarchical clustering methods, which allowed to distinguish three major clusters (Figures 4A and 4B). To confirm that these correspond to three different phases of the estrous cycle, we analyzed the expression of three known cycle dependent genes in the respective clusters, progesterone receptor (*Pgr*) and estrogen receptors (*Esr1* and *Gper1*). We found that these genes change indeed in the expected directions, both in the wildtype as well as the knockout animals (Figures 4C-E). Based on this finding, we performed the differential expression analysis on the three clusters separately. We found 21 differentially expressed genes in cluster 1 (DESeq2, adjusted P-Value ≤ 0.01; fold changes range from 0.75 to 1.59; Figure 3 – figure supplement 2), but still none for clusters 2 and 3. This suggests that *Udng3* acts mostly during the phase of high progesterone receptor and estrogen receptor 1 expression, and low G protein-coupled estrogen receptor 1 expression.

**Figure 4.**
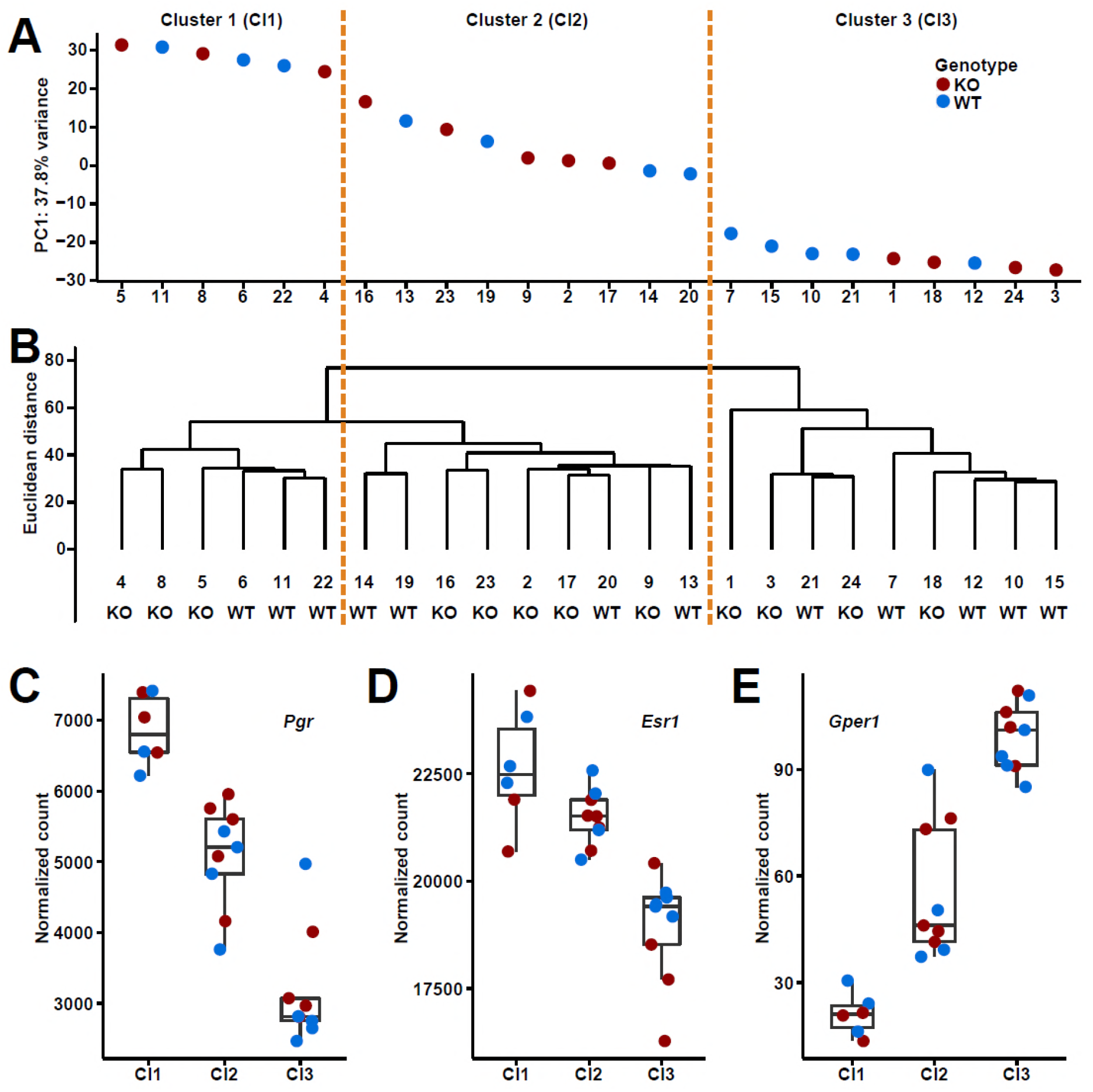
Clusters and expression levels in the 24 RNA-Seq samples of oviducts. (A) PC1 values from the PCA analysis, (B) hierarchical clustering result. Sample codes and genotypes are listed along X-axis. The 24 samples are assigned into three clusters accordingly. (C-E) The expression levels of three sex hormone receptor genes *(Pgr, Esr1, Gper1)* are shown by box plots. Figure 4 – figure supplement 1 shows the deletion patterns in the *Dcpp* gene region of the different populations (see text).

The top three differentially expressed genes belong all to a single young gene family, namely *Dcpp1* (ENSMUSG00000096445), *Dcpp2* (ENSMUSG00000096278) and *Dcpp3* (ENSMUSG00000057417), all three of which were significantly up-regulated in the knockout samples (DESeq2, fold changes: 1.45 for *Dcpp1,* 1.47 for *Dcpp2,* and 1.59 for *Dcpp3,* Figure 3 – figure supplement 2). These genes are specifically expressed in female and male reproductive organs and the thymus, and were previously found to function in oviducts to stimulate pre-implantation embryo development (Lee, Xu, Lee, & Yeung, 2006).

### Udng3 knockout phenotype

Given that the *Dcpp* genes are more highly expressed in *Udng3* knockouts, one could predict a higher implantation frequency of embryos, as it has been shown through experimental manipulation of *Dcpp* levels (Lee et al., 2006). We assessed the litters of pairs that were produced during our breeding experiments and found that the first litters from homozygous knockout females were produced after the same time as those from wildtype or heterozygous females (medians: 23 vs. 22 days; Table 3 - supplement 1), while we found that the second litters from homozygous knockout females were produced faster than those from wildtype or heterozygous females (medians: 23 vs. 38 days; Table 3 - supplement 1). To test this under more controlled conditions, we set up 10 mating pairs of homozygous knockout females with wildtype males and 10 wildtype pairs for control, all at approximately the same age at the start (8-9 weeks old). We found that the knockout and wildtype pairs had their first litter after the same time (medians: 23 vs. 22 days; Table 3 - supplement 1), while the knockout females had their second litter after a shorter time (medians: 24 vs. 36 days; Table 3 - supplement 1). Combining the total result from 36 mating pairs, we find that this difference is significant (two-tailed Wilcoxon rank sum test, P-Value = 0.042).

Interestingly, we found not only a timing difference for the second litter but also infanticide in about a quarter of the litters (4 out of 16) from homozygous females, but none in heterozygous or wildtype ones (two-tailed Fisher’s exact test, P-Value = 0.031; Table 3 - supplement 1). This could indicate that when the second litter follows too quickly, the females may be under strong postpartum stress resulting in partial killing of pups.

These results suggest that the loss of the *Udng3* gene should be detrimental to the animals in the wild.

Still, we see that the *M. m. domesticus* populations have secondarily lost this gene (Figure 2). Intriguingly, when inspecting the copy number variation data that we have produced previously (Pezer, Harr, Teschke, Babiker, & Tautz, 2015), we found that *Dcpp3* was also lost in *M. m. domesticus* populations (Figure 4 – figure supplement 1). Under the assumption that this results in an overall lowered expression of *Dcpp* RNAs, it could be considered to compensate for the loss of *Udng3.*

## Discussion

The aim of this study was to show that genes that have evolved only very recently out of previous non-coding regions can directly have a function, without further evolutionary adaptation. Out of a list of 119 candidate genes that have evolved *de novo* in the mouse lineage, we have chosen three more or less at random, only with the criterion to be particularly young and to represent different structural features and expression. We find that all three have an impact on the transcriptome and for all three we find traceable phenotypes related to their expression patterns when knocked out. Although the effects are subtle, at an evolutionary scale they can make a difference to the animals carrying them. Hence, we propose to give formal names to these genes. We name them after figures which emerged *de novo* as mythology characters in the Chinese classical novels *Journey to the West* and *Investiture of the Gods*, which were published in the 16th century. We name *Udng1* as *Sunwukong* (*Swk,* born from stone, *Journey to the West*), *Udng2* as *Leizhenzi* (*Lzhz*, born from thunderstorm, *Investiture of the Gods*), and *Udng3* as *Shiji* (*Shj,* born from stone, female, *Investiture of the Gods*).

### Functional de novo gene emergence

It has long been assumed that the emergence of function out of non-coding DNA regions must be rare, and if it occurs, the resulting genes would be far away from assuming a function. Our results do not support these assumptions. It is easy to find many well supported transcripts that could be considered to be true *de novo* genes. And three out of three chosen such genes can be shown to have functions. Hence, it would seem likely that most of the candidate genes in our curated list contribute aspects to the phenotype. Further, the fact that we neither observe patterns of ongoing positive selection, nor specifically accelerated evolution around these genes, suggests that they did not need additional adaptation to become functional. Although they have acquired a few additional substitutions, these are within the range of fixation of new neutral substitutions. This is in line with a similar analysis on a larger set of *de novo* ORFs in the mouse (Ruiz-Orera et al., 2018).

Our previous experiment with expressing random sequences in *E. coli* (Neme et al., 2017) had also suggested that the majority of them are not neutral, *i.e.,* they had an effect on the growth rates of the cells that carried them. We consider the question of whether this was a positive or negative effect as secondary (Tautz & Neme, 2018), since the evolutionary relevance is always in the context of other genes. This is best exemplified by *Udng3* / *Shj*. This has apparently a negative effect on the expression of its target genes. But through this negative effect, it provides apparently a life history advantage to the mice carrying it, since it suppresses too fast gestation that would otherwise have been caused by the duplicated genes. Thus, a negative effect results in a positive function in evolution.

We note also that an experiment that has expressed random peptides in plant (*Arabidopsis*) had a very high success rate of identifying associated phenotypes (Bao, Clancy, Carvalho, Elliott, & Folta, 2017).

One of the peptides that were functionally studied by these authors mediates an early flowering phenotype, which would self-evidently be a possible function for an ecological adaptation.

### Transcriptome changes and phenotypes

The fact that we see the disturbance of a whole transcriptomic network in the knockouts should of course not be interpreted to mean that the new genes interact directly with all of these other genes. We expect that even a single or a few interactions with other genes that are already part of a network could trigger this. Since our experimental design allowed a very high sensitivity to detect this, we were able to see the disturbance of many further interacting genes. We emphasize that the power of our analysis is much higher than in most transcriptomic studies, *i.e.,* we can see effects that would otherwise not be noted.

For *Udng2* / *Lzhz* the disturbed network has some functional coherence (extracellular matrix or cell motility functions), while the *Udng1* / *Swk* knockout results in rather broad effects. The fact that much fewer gene expression changes are seen for *Udng3* / *Shj* can be explained by the reduced power that we had in this experiment, due to the need to separate the data into three clusters. Similarly, the differences between females and males in the postnatal samples may be entirely due to different dispersions, rather than to sex-specific effects. But this question will need further study.

### Phenotype changes

None of the three knockout lines showed an overt phenotype, but we considered this also as *a priori* unlikely, given that a *de novo* evolved gene is expected to be only added to an existing network of genes. However, given the observed transcriptome changes, we were encouraged to apply a small set of phenotypic tests, relating to the respective major expression patterns of the genes. However, we consider the results from these tests only as preliminary at this stage. The behavioral tests in particular could be influenced by a variety of factors and would need repetition in much larger numbers. For example, the fact that “total distance” moved was measured in two behavioral tests (open field and novel object tests), but showed a significant difference in only one of the tests for *Udng1 / Swk* suggests a higher complexity. But at least the tendency was the same in both tests (shorter distance in knockouts). Still, we decided to not extend these tests for a larger number of *Udng2* mice.

For *Udng3 / Shj* we identified a possible direct link between the identified phenotype of a shorter gestation length in the knockouts and the transcriptomic changes. We found that the expression level of all three copies of *Dcpp* genes in C57BL/6N mice are enhanced in the *Udng3 /Shj* knockout animals. *Dcpp* expression is induced in the oviduct by pre-implantation embryos and is then secreted into the oviduct. This in turn stimulates the further maturation of the embryos and eventually the implantation (Lee et al., 2006). Hence, this is a system where a selfish tendency of embryos in expense of the resources of the mothers could develop. Accordingly, *Udng3 /Shj* could have found its function in controlling this expression. Intriguingly, the secondary loss of *Udng3 / Shj* in *M. m. domesticus* populations is accompanied by a loss of *Dccp3* in the same populations. This is compatible with the notion that an evolutionary conflict of interest exists for these interactions, whereby it remains open whether the loss of *Dcpp3* preceded the loss of *Udng3 / Shj* or vice versa.

### Conclusion

The notion that networks of gene interaction are far reaching and may have collective phenotypic effects has also been suggested in the context of quantitative trait genetics (Barton, Etheridge, & Veber, 2017; Boyle, Li, & Pritchard, 2017; Turelli, 2017). These authors have suggested that quantitative traits are eventually influenced by very many, if not all expressed genes. They emphasize also that modifying networks may be even more important than core networks in shaping quantitative phenotypes. Within the framework of such a concept, it is easy to see how a *de novo* evolved gene could integrate anywhere in the networks and lead to the subtle, but measurable perturbations on a whole set of genes, as shown in our data.

## Materials and Methods

### Ethics statement

The behavioral studies were approved by the supervising authority (Ministerium für Energiewende, Landwirtschaftliche Räume und Umwelt, Kiel) under the registration numbers V244-71173/2015, V244-4415/2017 and V244-47238/17. Animals were kept according to FELASA (Federation of European Laboratory Animal Science Association) guidelines, with the permit from the Veterinäramt Kreis Plön: 1401-144/PLÖ-004697. The respective animal welfare officer at the University of Kiel was informed about the sacrifice of the animals for this study.

### Genome-wide identification of de novo genes

We modified previous phylostratigraphy and synteny-based methods to identify *Mus*-specific *de novo* protein-coding genes from intergenic regions. We started with mouse proteins annotated in Ensembl (Version 80) (Zerbino et al., 2018) (1) with protein length not smaller than 30 amino acids, (2) with a start codon at the beginning of the ORF, (3) with a stop codon at the end of the ORF, (4) without stop codons within the annotated ORF. For the phylostratigraphy-based strategy, in order to save computational time, we first used NCBI BLASTP (2.5.0+) to align low complexity region masked mouse protein sequences to rat protein sequences annotated in Ensembl (Version 80) and filtered out the mouse sequences having hits with E-values smaller than 1 × 10^−7^. This removes all conserved genes. Next we used NCBI BLASTP (2.5.0+) to align the remaining low complexity region masked sequences to NCBI nr protein sequences (10 Nov. 2016) (O’Leary et al., 2016) and filtered out the mouse sequences having non-genus *Mus* hits with E-values smaller than 1×10^−3^ according to (Neme & Tautz, 2013). The genes remaining after these filtering steps are the candidates for the *de novo* evolved genes. In order to deal also with proteins having low complexity regions, we further applied a synteny-based strategy on the rest proteins by taking advantage of the Chain annotation from Comparative Genomics of UCSC Genome Browser (“http://genome.ucsc.edu/”) (Kent et al., 2002). We filtered out the proteins encoded on unassembled scaffolds because their chromosome information is not compatible between Ensembl and UCSC annotations. We only compared rat and human proteins with mouse proteins because their genomes are well assembled and genes are well annotated. We performed the same procedures on rat and human data separately, and used “mm10.rn5.all.chain” and “rn5ToRn6.over.chain” from UCSC and gene annotation from Ensembl (Version 80) for rat, and “mm10.hg38.all.chain” from UCSC and gene annotation from Ensembl (Version 80) for human. For each mouse gene, if its ORF overlaps with any ORFs in the rat or human mapping regions in Chain annotation, we aligned its protein sequence to those protein sequences with program water from EMBOSS (6.5.7.0) (Rice, Longden, & Bleasby, 2000); if one of the alignment scores is not smaller than 40, we filtered out the protein. The remaining 119 genes are the candidates for the following analysis and the pool for us to select genes for detailed functional experiments.

### ENCODE RNA-Seq analysis

We downloaded the raw read files of 135 strand-specific paired-end RNA-Seq samples generated by the lab of Thomas Gingeras, CSHL from ENCODE (Consortium, 2012; Sloan et al., 2016) including 35 tissues from different organs and different developmental stages, and each of them had multiple biological or technical replicates (see list in Figure 1 – figure supplement 2). We trimmed the raw reads with Trimmomatic (0.35) (Bolger, Lohse, & Usadel, 2014), and only used paired-end reads left for the following analyses. We mapped the trimmed reads to the mouse genome GRCm38 (Mouse Genome Sequencing et al., 2002; Zerbino et al., 2018) with HISAT2 (2.0.4) (Kim, Langmead, & Salzberg, 2015) and SAMtools (1.3.1) (H. Li et al., 2009), and took advantage of the mouse gene annotation in Ensembl (Version 80) by using the --ss and --exon options of hisat2-build. We assembled transcripts in each sample, and merged annotated transcripts in Ensembl (Version 80) and all assembled transcripts with StringTie (1.3.4d) (Pertea et al., 2015). Then we estimated the abundances of transcripts, FPKM values, in each sample with StringTie (1.3.4d). For each tissue, we summarized the FPKM values of each transcript by averaging the values from multiple biological or technical replicates; and if a gene has multiple transcripts, we assigned the summary of the FPKM values of the transcripts as the transcriptional abundance of the gene.

### Ribosome profiling and proteomics analysis

We downloaded the datasets that included both strand-specific ribosome profiling (Ribo-Seq) and RNA-Seq experiments of the same mouse samples from Gene Expression Omnibus (Barrett et al., 2013) under accession numbers GSE51424 (Gonzalez et al., 2014), GSE72064 (Cho et al., 2015), GSE41426 (Djiane et al., 2013), GSE22001 (Guo, Ingolia, Weissman, & Bartel, 2010), GSE62134 (Diaz-Munoz et al., 2015), and GSE50983 (Castaneda et al., 2014), which corresponded to brain, hippocampus, neural ES cells, heart, skeletal muscle, neutrophils, splenic B cells, and testis. Ribo-seq datasets were depleted of possible rRNA contaminants by discarding reads mapped to annotated rRNAs, and then the rest reads were mapped to GRCm38 (Mouse Genome Sequencing et al., 2002; Zerbino et al., 2018) with Bowtie2 (2.1.0) (Langmead & Salzberg, 2012). RNA-Seq reads were mapped to the mouse genome GRCm38 with TopHat2 (2.0.8) (Kim et al., 2013). Then we applied RiboTaper (1.3) (Calviello et al., 2016) which used the triplet periodicity of ribosomal footprints to identify translated regions to the bam files. Mouse GENCODE Gene Set M5 (Ensembl Version 80) (Mudge & Harrow, 2015) was used as gene annotation input. The Ribo-seq read lengths to use and the distance cutoffs to define the positions of P-sites were determined from the metaplots around annotated start and stop codons as shown below.

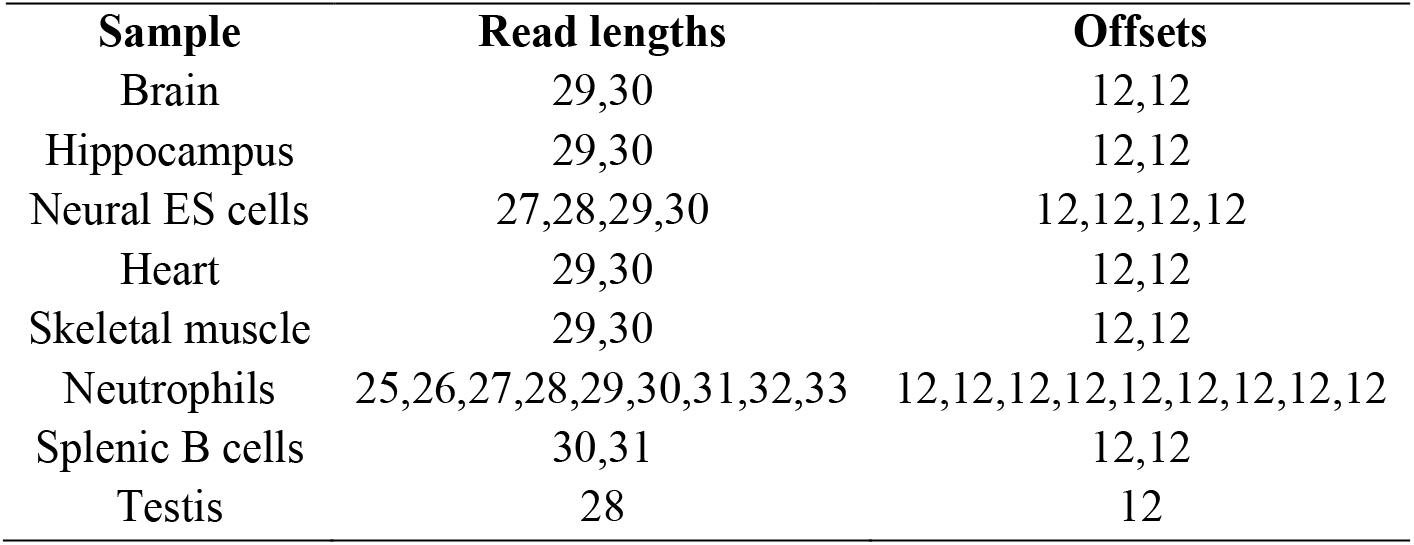

All mouse peptide evidence from large-scale mass spectrometry studies was retrieved from PRIDE (09 Aug. 2015) (Vizcaino et al., 2016) and PeptideAtlas (31 Jul. 2015) (Desiere et al., 2006) databases. We performed the same procedures on PRIDE and PeptideAtlas data separately following the method described in (Xie et al., 2012). In brief, if the whole sequence of a peptide was identical to one fragment of the tested *de novo* protein sequence, and had at least two amino acids difference compared to all the fragments of other protein sequences in the mouse genome, the peptide was considered to be convincing evidence for the translational expression of the respective *de novo* protein.

### Molecular patterns of de novo genes

The exon number of a gene was assigned as the exon number of the transcript having highest FPKM value among all the transcripts of the gene. The intrinsic structural disorder of proteins was predicted using IUPred (Dosztanyi, Csizmok, Tompa, & Simon, 2005), long prediction type was used. The intrinsic structural disorder score of a protein was assigned as the average of the scores of all its amino acids. The hydrophobic clusters of proteins were predicted using SEG-HCA (Faure & Callebaut, 2013), and then the fraction of the sequence covered by hydrophobic clusters for each protein was calculated.

### RT-PCR

The ovaries, oviducts, uterus, and gonadal fat pad from wildtype *Udng3* females were carefully collected and immediately frozen in liquid nitrogen. Total RNAs from those tissues were purified using QIAGEN RNeasy Microarray Tissue Mini Kit (Catalog no. 73304), and the genomic DNAs were removed using DNase I, RNase-free (Catalog no. 74106). The first strand cDNAs were synthesized using the Thermo Scientific RevertAid First Strand cDNA Synthesis Kit (Catalog no. K1622) by targeting poly-A mRNAs with oligo dT primers. Two pairs of primers targeted on the two junctions of the *Udng3* gene structure and a pair of primers targeted on a control gene *Uba1* were used. The sequences of the primers are shown below. PCR was done under standard conditions for 38 cycles.

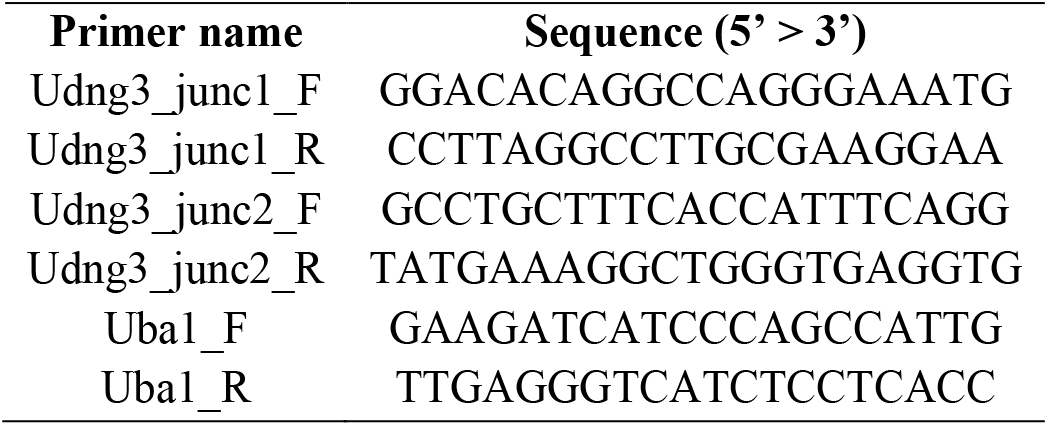

### Genomic sequences of Udng1, Udng2, and Udng3 loci from wild mice

The genomic sequences from *M. spretus* (8 individuals), *M. m. castaneus* (10 individuals), *M. m. musculus* from Kazakhstan (8 individuals), *M. m. musculus* from Afghanistan (6 individuals), *M. m. musculus* from Czech Republic (8 individuals), *M. m. domesticus* from Iran (8 individuals), *M. m. domesticus* from Germany (11 individuals), and *M. m. domesticus* from France (8 individuals) were retrieved from the whole genome sequencing data in (Harr et al., 2016).

The genomic sequences from *A. uralensis* (4 individuals), *M. mattheyi* (4 individuals), and *M. spicilegus* (4 individuals) were determined by Sanger sequencing of the PCR fragments from the genomic DNAs purified with salt precipitation. The PCR primers listed below were designed according to the whole genome sequencing data of the three species in (Neme & Tautz, 2016). There were only few reads from the *A. uralensis* whole genome sequencing data mapped to the *Udng3* locus in the reference genome, and we did not design primers to try to determine the sequences, because the *A. uralensis* genomic sequence at this locus is very different from the reference *(M. m. domesticus),* and the *Udng3* ORF does not exist.

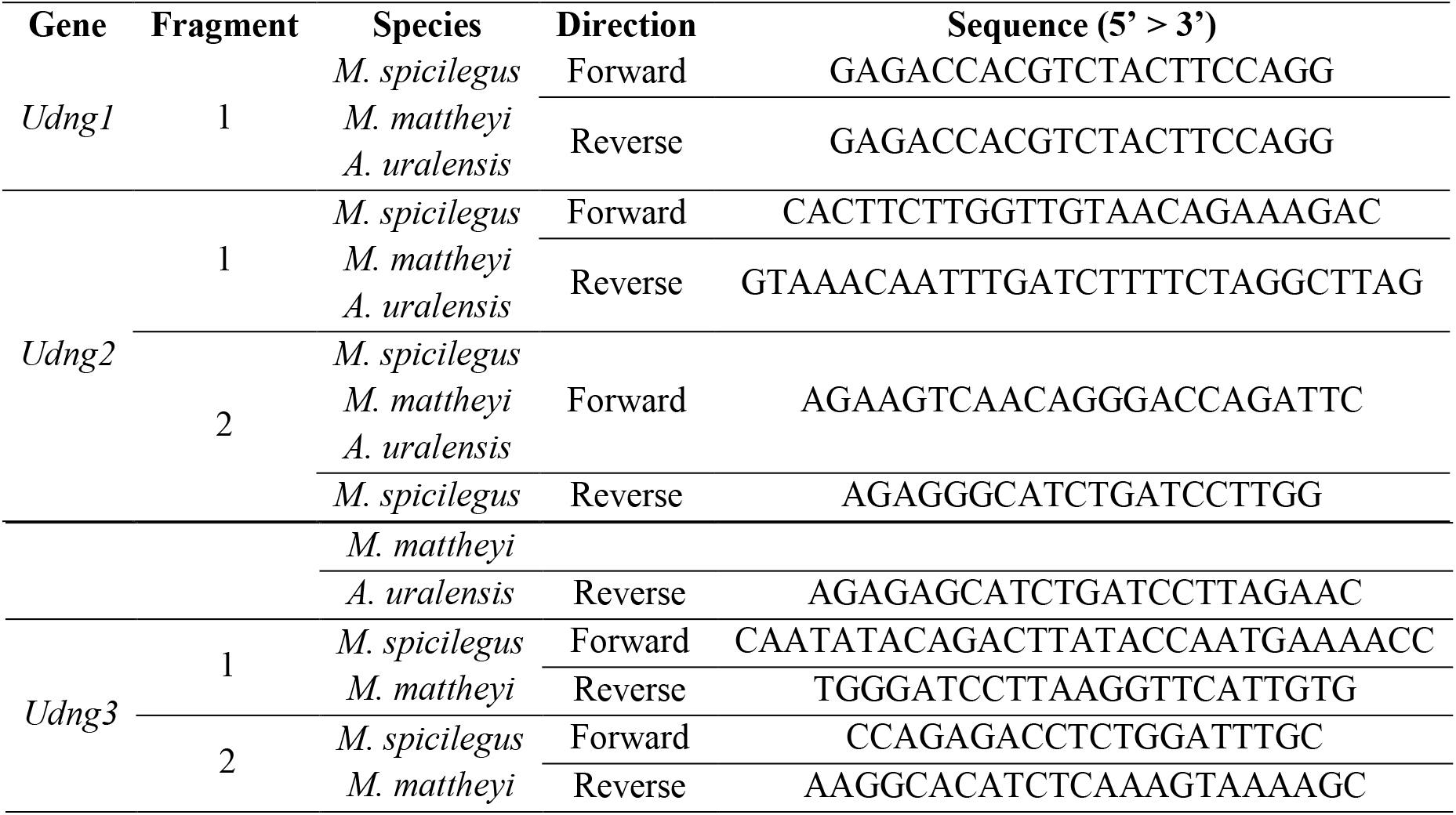

### Phylogenetic distance analysis

Whole genome sequencing data in (Harr et al., 2016) and (Neme & Tautz, 2016) were used to obtain the average distances for the taxa in this analysis. For each individual, the mean mapping coverage was calculated using ANGSD (0.921-10-g2d8881c) (Korneliussen, Albrechtsen, & Nielsen, 2014) with the options “-doDepth 1 -doCounts 1 -minQ 20 -minMapQ 30 -maxDepth 99999”. Then, ANGSD (0.921-10-g2d8881c) was used to extract the consensus sequence for each population accounting for the number of individuals and the average mapping coverage per population (mean + 3 times standard deviation) with the options “-doFasta 2 -doCounts 1 -maxDepth 99999 -minQ 20 -minMapQ 30 -minIndDepth 5 – setMinDepthInd 5 -minInd X1 -setMinDepth X2 -setMaxDepthInd X3 -setMaxDepth X4”. X1, X2, X3, and X4 are listed below. The consensus sequences of the mouse populations were used to calculate the Jukes-Cantor distances for 10,000 random non-overlapping 25 kbp windows from the autosomes with APE (5.1, “dist.dna” function) (Paradis, Claude, & Strimmer, 2004). The average distances obtained in this way are provided in Figure 2 – figure supplement 2. The expected distances for the three genes in Table 2 were calculated by multiplying the length of the gap-free alignment with the average distances. The observed values were retrieved from the distance table of the alignments using Geneious (11.1.2).

| Population | Mean coverage | Standard deviation of coverage | X1 | X2 | X3 | X4 |
| --- | --- | --- | --- | --- | --- | --- |
| *A. uralensis* | 17.912 | 23.999 | 1 | 5 | 90 | 90 |
| *M. mattheyi* | 23.304 | 83.028 | 1 | 5 | 273 | 273 |
| *M. spicilegus* | 25.138 | 24.627 | 1 | 5 | 100 | 100 |
| *M. spretus* | 24.885 | 14.216 | 4 | 20 | 68 | 54 |
| *M. m. castaneus* | 14.015 | 7.573 | 5 | 25 | 37 | 370 |
| *M. m. musculus* from Afghanistan | 17.768 | 58.551 | 3 | 15 | 59 | 354 |
| *M. m. musculus* from Kazakhstan | 25.123 | 15.975 | 4 | 20 | 74 | 592 |
| *M. m. musculus* from Czech Republic | 24.338 | 14.103 | 4 | 20 | 67 | 536 |
| *M. m. domesticus* from Iran | 20.249 | 9.820 | 4 | 20 | 50 | 400 |
| *M. m. domesticus* from Germany | 21.639 | 10.518 | 4 | 20 | 54 | 432 |
| *M. m. domesticus* from France | 21.499 | 10.027 | 4 | 20 | 52 | 416 |

### Mouse knockout lines

The line with allele *A930004D18Rik^tm1a(EUCOMM)Wtsi^* (genetic background: C57BL/6N) was obtained from the European Mouse Mutant Archive (EMMA). We converted it to the *Udng1* knockout line *(tm1b)* using a cell-permeable Cre recombinase in order to delete the coding exon together with the selection cassette according to the method described in (Ryder et al., 2014). In brief, the females from the line were super-ovulated and were then mated with the males from the line. The 2-cell embryos were collected and treated with HTN-Cre from Excellgen (Catalog no. RP-7). Then they were transferred into 0.5-day pseudo-pregnant females. The alleles of the pups were confirmed by PCR and Sanger sequencing, and only the mice with black coat color were used for further breeding and experiments.

The knockout line for *Udng2* with allele *A830005F24Rik^tm1.1(KOMP)Mbp^* (genetic background: C57BL/6N) was obtained from the Knock-Out Mouse Project (KOMP).

*Udng3* was also originally targeted by KOMP, but the line was lost. Hence, we obtained a custom-made CRISPR/Cas9 line from the Mouse Biology Program (MBP). The guide RNA was designed to target the beginning of the ORF in the second coding exon and away from the splicing site (genomic DNA target: 5 ‘ TGCTCCATCTGCTTTTCAGG 3’). We obtained three mosaic frameshift knockout mice (genetic background: C57BL/6N). Then we mated them with the wildtypes from the same litters to have heterozygous pups, and selected one female and one male with a heterozygous 7-bp deletion as the founding pair for further breeding and experiments.

Primers for genotyping the three lines are listed below.

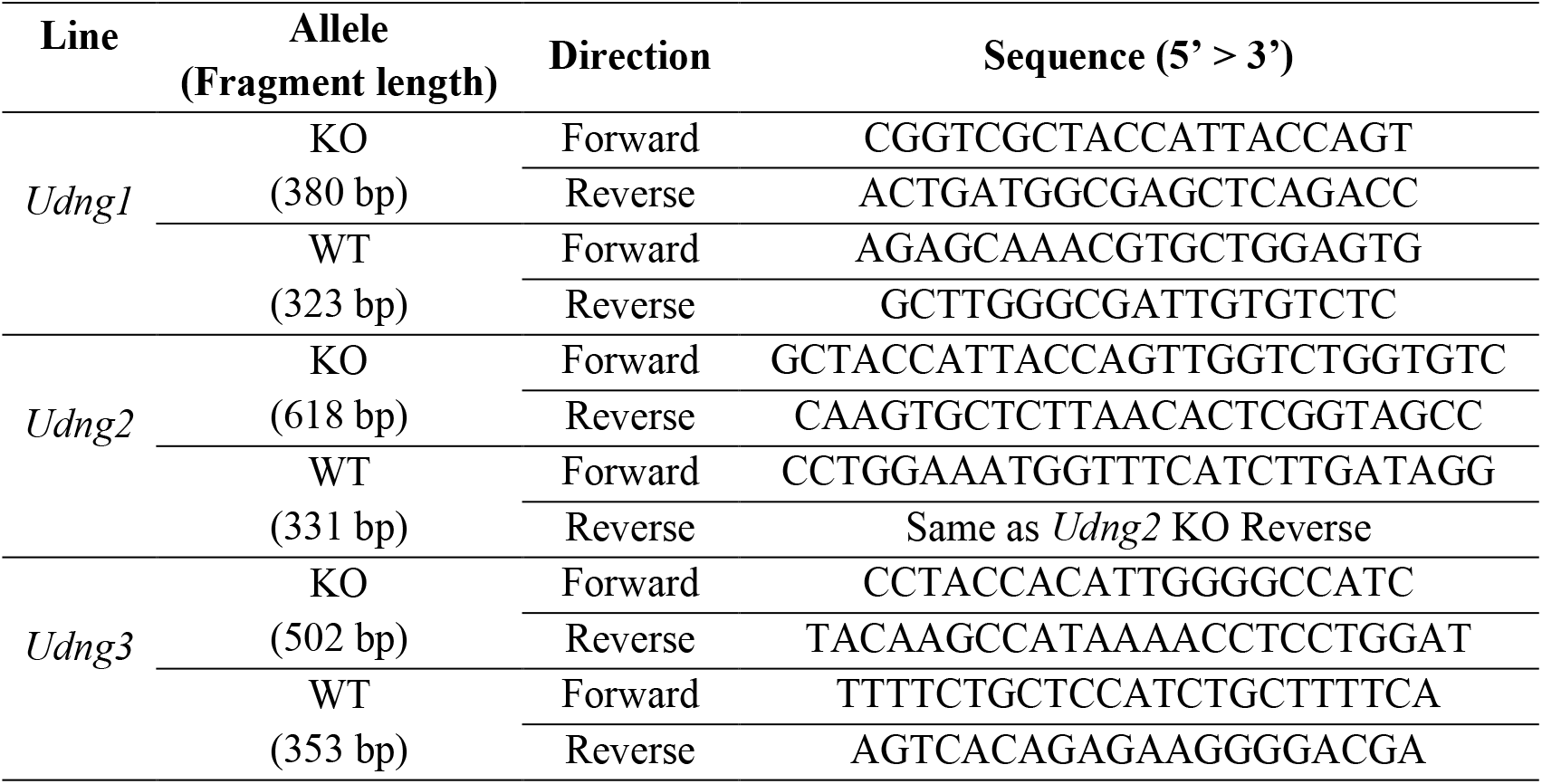

### Power analysis for RNA-Seq

RnaSeqSampleSize (1.6.0) (Zhao, Li, Guo, Sheng, & Shyr, 2018) was used for power analysis. Specifically, we used the est_power function, and set parameters w (ratio of normalization factors between two groups) as 1, alpha (significance level) as 3.3 × 10^−6^. Then we traversed all 98,670 possible combinations of N (sample size) from 3 to 13, rho (fold change) from 1.05 to 1.5, lambda0 (read count) from 4 to 65,536, and phi0 (dispersion) from 0.00025 to 1.024 to calculate the power values.

To calculate the power of each gene in each of our real RNA-Seq datasets, we also used the est_power function with the parameters w as 1, alpha as 3.3 × 10^−6^, and n as the real sample size, and rho (2_log2FoldChange|_), lambda0 (baseMean), and phi0 (dispersion) estimated by DESeq2 (1.14.1) (Love, Huber, & Anders, 2014) based on the real data.

### lacZ overexpression

Primary mouse embryonic fibroblasts (MEFs) used for overexpression were obtained from C57BL/6 mice. Specifically, we dissected 13.5-14.5 dpc embryos from uteruses and extraembryonic membranes into PBS (Lonza, Catalog no. BE17-512F); discarded heads and soft tissues and washed the carcasses with PBS; cut the carcasses into 2-3 mm pieces; transferred them into 50 ml Falcon tubes and added 5-20 ml

Trypsin-EDTA (Gibco, Catalog no. 25300-054); vortexed and incubated for 10 minutes at 37°C; vortexed again and incubated for 10 minutes at 37°C; inactivated trypsin by adding 2 volumes of medium (500 ml DMEM (Lonza, Catalog no. BE12-733F), 55 ml FBS (PAN, Catalog no. P30-3702), 5.5 ml glutamine (Lonza, Catalog no. BE17-605E), 5.5 ml penicillin (5,000 U/ml) / streptomycin (5,000 μg/ml) (Lonza, Catalog no. DE17-603)); pipetted up and down to get single cell suspension; plated cells and incubated overnight.

We separately cloned the fragment of the *lacZ* ORF from the *Udng2* knockout allele *(Udng1* and *Udng2* knockout alleles have the identical *lacZ* ORF) and its reverse complement fragment into pVITRO2-neo-GFP/LacZ expression vector (Catalog no. pvitro2-ngfplacz) to replace its own *lacZ* ORF using homologous recombination method, and then purified the plasmids with QIAGEN EndoFree Plasmid Maxi Kit (Catalog no. 12362). The replacements in the vectors were confirmed by PCR and Sanger sequencing. Ten independent transfections for each of the two plasmids into the P2 MEFs were performed separately with Amaxa Mouse/Rat Hepatocyte Nucleofector™ Kit (Catalog no. VPL-1004) according to manufacturer’s recommendation. Transfected cells were grown in the medium (see above). Cells were incubated at 37°C in 5% CO_2_ atmosphere as a pH regulator. The expression of *lacZ* in *lacZ* overexpressed cells but not in reverse *lacZ* overexpressed cells was confirmed using a β-Galactosidase Staining Kit (Catalog no. K802-250). Total RNAs from the transfected cells were purified using QIAGEN RNeasy Mini Kit (Catalog no. 74106) 48 hours after transfection.

### RNA-Seq and data analysis

The heads of postnatal 0.5-day *Udng1* and *Udng2* pups, the 12.5-day *Udng1* embryos, and the oviducts of 10-11 weeks old *Udng3* females were carefully collected and immediately frozen with liquid nitrogen. Then, for all these samples, total RNAs were purified using QIAGEN RNeasy Microarray Tissue Mini Kit (Catalog no. 73304). All RNA samples, including the total RNAs purified from the transfected MEF cells, were prepared using Illumina TruSeq Stranded mRNA HT Library Prep Kit (Catalog no. RS-122-2103), and sequenced using Illumina NextSeq 500 and NextSeq 500/550 High Output v2 Kit (150 cycles) (Catalog no. FC-404-2002). All procedures were performed in standardized and parallel way.

Raw sequencing outputs were converted to FASTQ files with bcl2fastq (2.17.1.14), and reads were trimmed with Trimmomatic (0.35) (Bolger et al., 2014). Only paired-end reads left were used for following analyses. We mapped the trimmed reads to mouse genome GRCm38 (Mouse Genome Sequencing et al., 2002; Zerbino et al., 2018) with HISAT2 (2.0.4) (Kim et al., 2015) and SAMtools (1.3.1) (H. Li & Durbin, 2009), and took advantage of the mouse gene annotation in Ensembl (Version 86) by using the --ss and --exon options of hisat2-build. We counted fragments mapped to the genes annotated by Ensembl (Version 86) with HTSeq (0.6.1p1) (Anders, Pyl, & Huber, 2015), and performed differential expression analysis with DESeq2 (1.14.1) (Love et al., 2014). Besides the DESeq2 default outputs, we also added the dispersions estimated by DESeq2 (1.14.1) and the powers calculated by RnaSeqSampleSize (1.6.0) (Zhao et al., 2018) (see *Power analysis for RNA-Seq)* into the outputs.

KOBAS (2.0) (Xie et al., 2011) was used for functional enrichment analysis.

For the RNA-Seq of the oviducts of *Udng3* females, principle component analysis and hierarchical clustering with Euclidean distance and complete agglomeration method on the variance stabilized transformed fragment counts were also performed using DESeq2 (1.14.1) to assign the 24 samples into three clusters.

### Whole genome sequencing of the Udng3 founding pair and off-target analysis

The genomic DNAs from the founding pair were purified with salt precipitation. Then the samples were prepared with Illumina TruSeq Nano DNA HT Library Prep Kit (Catalog no. FC-121-4003), and sequenced on HiSeq 2500 with TruSeq PE Cluster Kit v3-cBot-HS (Catalog no. PE-401-3001) and HiSeq Rapid SBS Kit v2 (500 cycles) (Catalog no. FC-402-4023). The reads were 2 × 250 bp in order to have good power to detect indels.

We followed GATK Best Practices (Van der Auwera et al., 2013) to call variants. Specifically, we mapped the reads to mouse genome GRCm38 (Mouse Genome Sequencing et al., 2002; Zerbino et al., 2018) with BWA (0.7.15-r1140) (H. Li & Durbin, 2009), and marked duplicates with Picard (2.9.0) (http://broadinstitute.github.io/picard), and realigned around the indels founded in C57BL/6NJ line (Keane et al., 2011) with GATK (3.7), and recalibrated base quality scores with GATK (3.7) using variants founded in C57BL/6NJ line (Keane et al., 2011) to get analysis-ready reads. We assessed coverage with GATK (3.7) and SAMtools (1.3.1) (H. Li et al., 2009), and the coverage of female was 35.48 X and the one of male was 35.09 X. High coverages also provided good power to detect indels. We called variants with GATK (3.7), and applied generic hard filters with GATK (3.7): “QD < 2.0 ∥ FS > 60.0 ∥ MQ < 40.0 ∥ MQRankSum < −12.5 ∥ ReadPosRankSum < −8.0 ∥ SOR > 3.0” for SNVs and “QD < 2.0 ∥ FS > 200.0 ∥ ReadPosRankSum < −20.0 ∥ SOR > 10.0” for indels. We found 80375 SNVs and 73387 indels in the female and 81213 SNVs and 71857 indels in the male.

347 potential off-target sites were predicted on “http://crispr.mit.edu:8079/” based on mouse genome mm9. 343 of them still existed in mouse genome mm10 after converting by liftOver (26 Jan. 2015) (Kent et al., 2002), and the four missing sites were rank low anyway: 131, 132, 143, and 200. GATK (3.7) was used to look for variants found in the whole genome sequencing in the 100 bp regions around the 343 sites. In addition, the reads mapped to the regions around the top 20 sites were manually checked in both samples.

### Behavioral tests

The following behavioral tests were performed on the *Udng1* and *Udng2* mice used in this study: elevated plus maze test, open field test and novel object test. All tests were recorded on video using a VK-13165 Eneo camera mounted directly above the experimental set-up and behaviors were measured using VideoMot2 (TSE Systems). All tests were filmed in the same room under similar lighting conditions (less than 200 lux). All lights faced the ceiling in order to avoid any glare or reflections within the test arenas. For the elevated plus maze we used an arena that was designed for testing wild mice. It was constructed as two perpendicular arms using PVC plastic and acrylic glass, and was 80 cm above ground. The dark arms of the maze were made with grey PVC plastic sides, with a white PVC plastic bottom. The dark arms were 50 cm long, 10 cm wide and 40 cm deep. Open arms had same dimensions, except that the walls were made of acrylic glass instead of grey plastic. For testing, each mouse was placed at the center of the arena at the beginning of the test using a transparent plastic transfer pipe. Mice were filmed inside the test arena for 5 minutes (Holmes, Parmigiani, Ferrari, Palanza, & Rodgers, 2000). VideoMot2 (TSE Systems) was used to measure the time which the mouse spent in the dark arm, the light arm, and the center of the maze. After each experiment, the test arena was cleaned with 30% ethanol.

The open field arena was made of white PVC plastic and measured 60 x 60 cm, and the walls were 60 cm high. The arena was placed directly beneath a security camera and measurements were taken using VideoMot2 (TSE Systems). At the beginning of the experiment, the mouse was placed at the center of the arena using a transparent plastic transfer pipe. Each mouse was filmed for 5 minutes. Measurements taken during the open field test included the amount of time spent at the wall of the arena (up to 8 cm away from the wall) and the distance travel during the experiment (Yuen, Pillay, Heinrichs, Schoepf, & Schradin, 2016). After each experiment, the test arena was cleaned with 30% ethanol.

The novel object test was carried out in the same arena as the open field test. The arena was placed directly beneath a security camera and measurements were taken using VideoMot2 (TSE Systems). At the beginning of the experiment, the mouse was placed at the center of the arena using a transparent plastic transfer pipe along with a toy made of colored building blocks (Lego). Each mouse was filmed for 5 minutes. Measurements taken during the novel object test included the latency to investigate the novel object, the number of visits to the novel object, and the distance travel during the experiment. The number of visits to the novel object was accessed based on visits to an area of 7.5 cm around the novel object (Yuen et al., 2016). After each experiment, the test arena and novel object were cleaned with 30% ethanol. All the measured *Udng1* and *Udng2* mice are adult males. They were genotyped in advance, matched between knockouts and wildtypes. The genotypes were then masked to the experimenter. Their ages were from 11 to 17 weeks old for *Udng1* and from 15 to 25 weeks old for *Udng2.* Each mouse stayed alone in the cage in a room with only male mice at least two weeks before measurements. 40 *Udng1* mice measured by elevated plus maze test, open field test, and novel object test were divided into two groups (20 in Group A and 20 in Group B) and were measured in two different days for the same test. 36 *Udng2* mice measured by elevated plus maze test were divided into three groups (12 in Group A, 8 in Group B, and 16 in Group C) and were measured in three different days. For the open field test and novel object test, only the group A mice could be measured. The order of the mice to be measured in each group was randomly shuffled.

Nested ranks test (Thompson, Smouse, Scofield, & Sork, 2014) was used for the statistical analyses to compare the parameters in each behavioral tests between knockouts and wildtypes. It is a non-parametric linear mixed model test, and uses the genotype as the fixed effect and the group membership as the random effect. For the parameters of the behavioral tests having only one group, it is essentially identical to one-tailed Wilcoxon rank sum test.

### Limb morphology

Mouse limbs were scanned using a computer tomograph (micro-CT-vivaCT 40, Scanco, Bruettisellen, Switzerland; energy: 70 kVp, intensity: 114 μA, voxelsize: 38 μm). Further, three-dimensional cross-sections were generated with a resolution of one cross-section per 0.038 mm. Two 3D landmarks were located at the endpoints of each limb bone using the TINA landmarking tool (Schunke, Bromiley, Tautz, & Thacker, 2012), and the linear distance between the two landmarks were calculated for statistical analyses. Detailed description of landmarks for each bone was previously reported in (Skrabar et al., 2018). Measurements were obtained from the right side of three forelimb bones (humerus, ulna, and metacarpal bone) and three hindlimb bones (femur, tibia, and metatarsal bone).

40 *Udng1* adult males at an age between 13-19 weeks were measured. They were genotyped in advance, matched between knockouts and wildtypes and then the genotypes were masked to the experimenter. The order of the mice to be measured in each group was randomly shuffled.

### Fertility test

*Udng3* mating pairs were set up for the fertility test. The female and male in each pair were 8-9 weeks old when the mating was started. All the males were wildtype, and 10 females were homozygous knockout and the other 10 were wildtype. The time (days) having the 1st or 2nd litters, the numbers of pups of the 1st or 2nd litters, and whether the pups were eaten later for each mating pair were carefully observed and recorded by animal caretakers who were blind about the genotypes.

### Data availability

The ENA BioProject accession number for the sequencing data reported in this study is PRJEB28348.

## Acknowledgments

The authors are grateful to R. Neme for generating a first version of the candidate gene list; J. Ruiz-Orera for generating the bam files from Ribo-Seq datasets; C. Pfeifle, A. Vock, A. Jonas, C. Medina, and S. Holz for keeping the mice used in this project; S. von Merten, C. Pfeifle, H. Harre, and W. Rasmus for helping mouse phenotyping experiments; B. Kleinhenz for helping cell culture experiments; E. Blohm-Sievers for helping mouse genotyping and Sanger sequencing; C. Burghardt, E. McConnell, and H. Buhtz for helping Illumina sequencing. The mouse line with *Udng1* targeted allele used for this project was obtained from EMMA; the *Udng2* knockout mouse line used for this project was obtained from KOMP; and the *Udng3* knockout mouse line used for this project was obtained from MBP. This work was supported by a European Research Council advanced grant to DT (NewGenes – 322564).

---

**Figure 2 – figure supplement 1.**
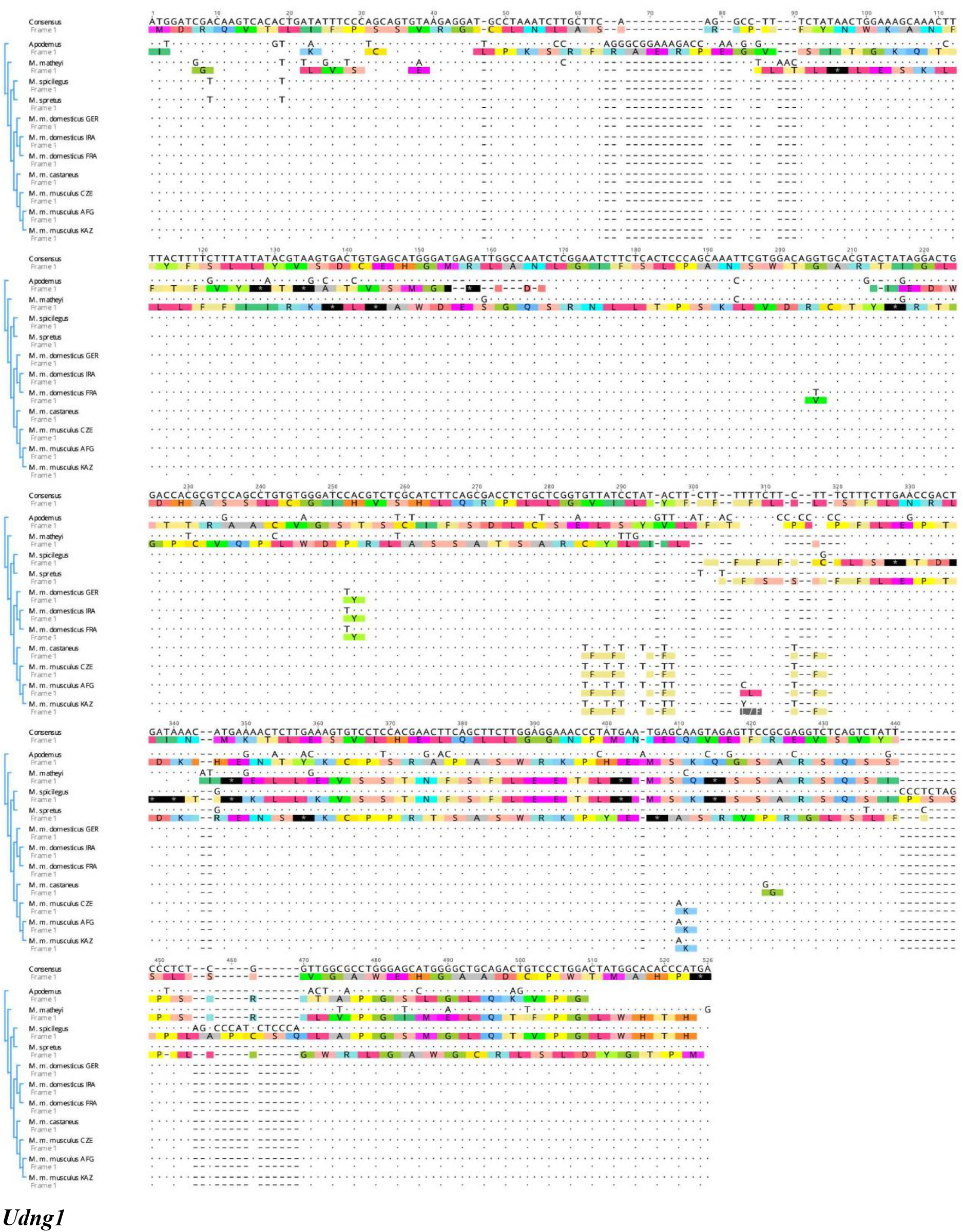
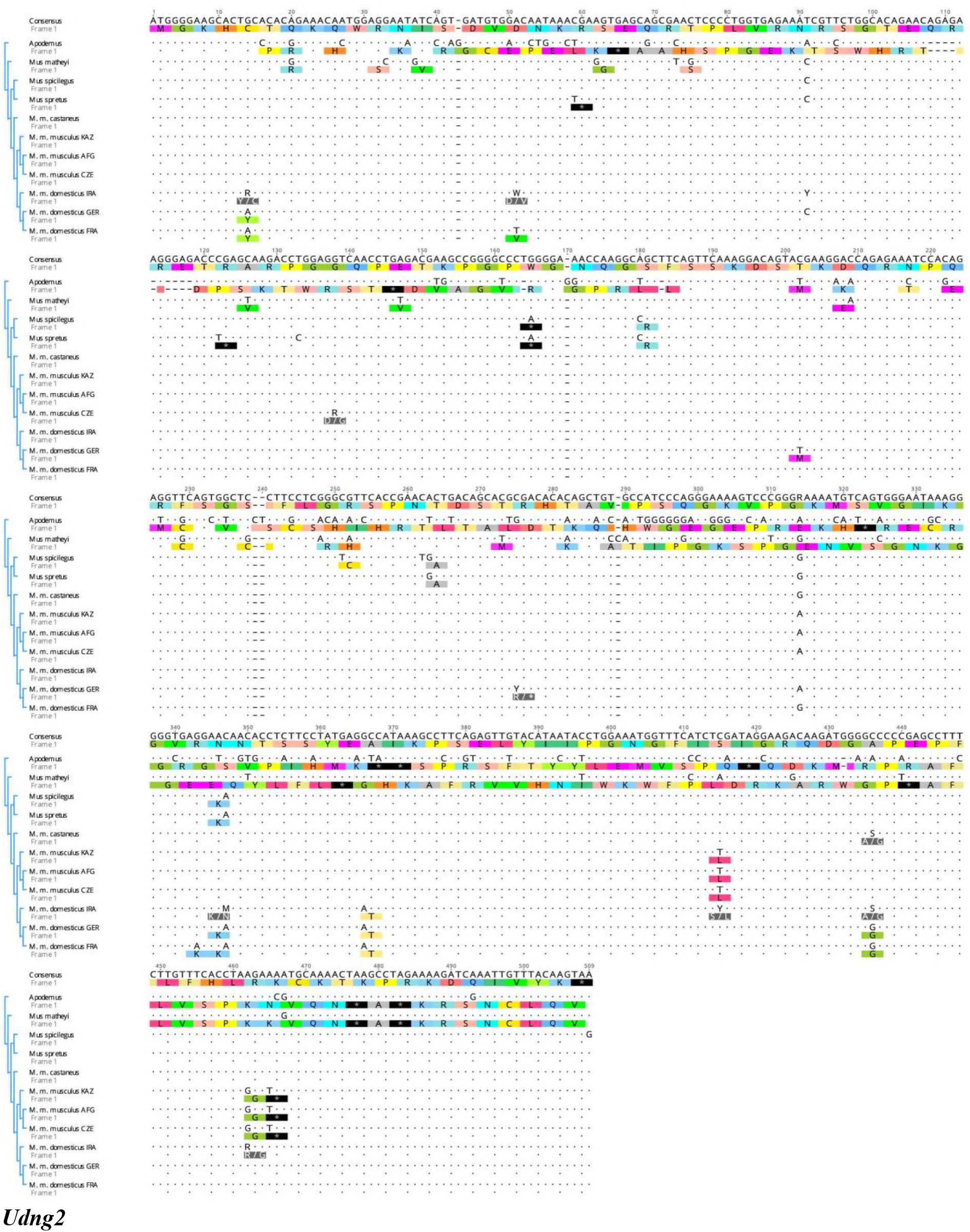
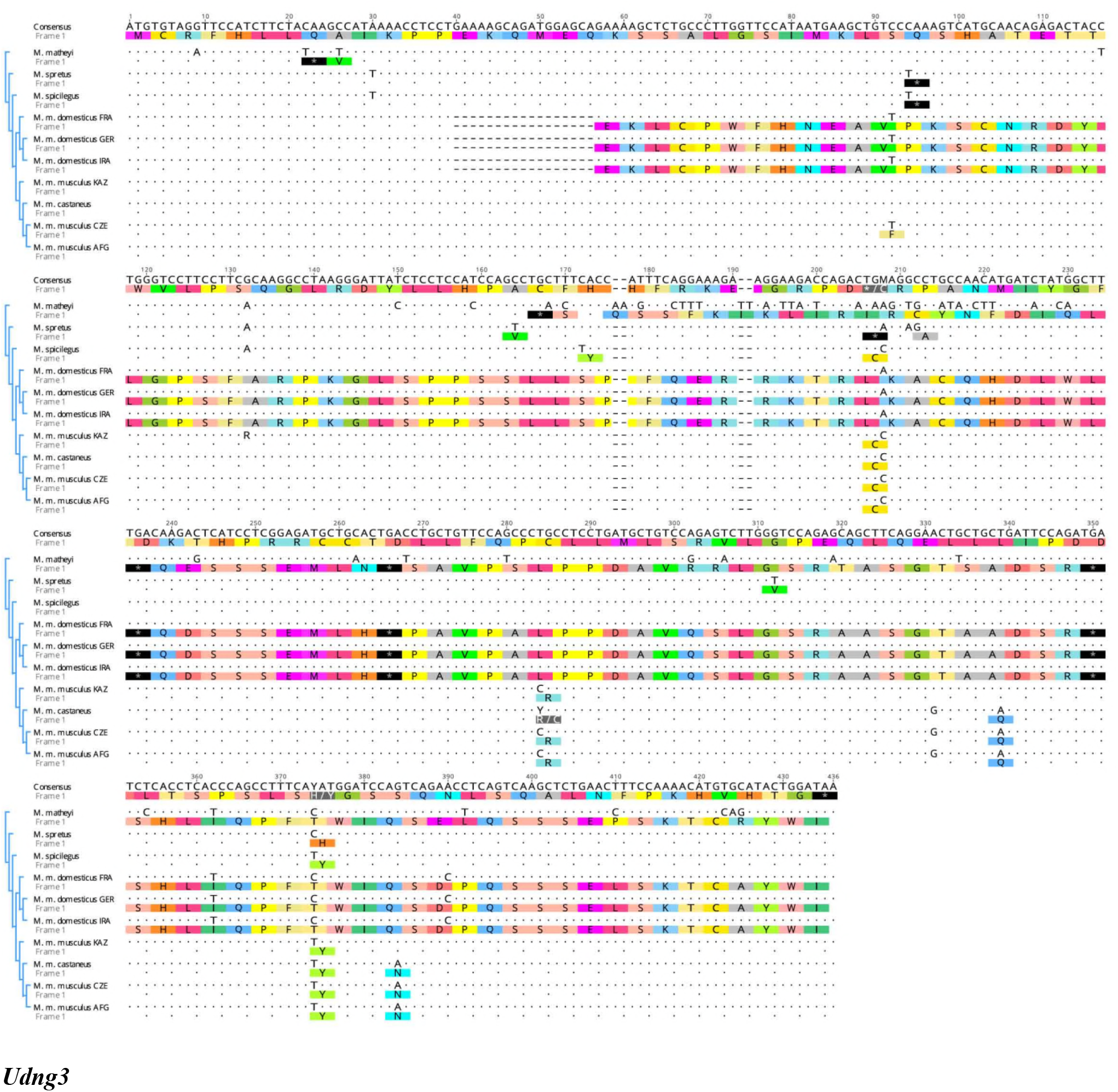
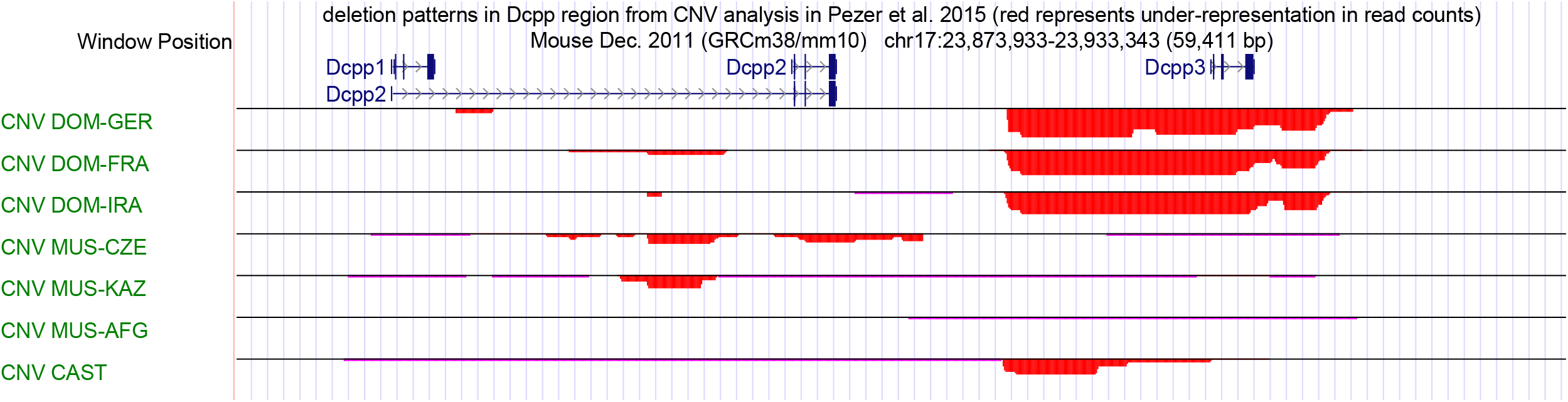
The alignments of *Udng1, Udng2* and *Udng3* ORF sequences from mouse species (*Mus =* M.), subspecies (*Mus musculus* = *M. m.)* and the outgroup *Apodemus.* Three populations each are represented for *M. m. musculus* (KAZ = from Kazakhstan, AFG = from Afghanistan, CZE = from Czech Republic) and *M. m. domesticus* (IRA = from Iran, FRA = from France, GER = from Germany). All sequences are aligned to a consensus sequence that is produced as a consensus across all sequences shown. Identical positions are marked by a dot, replacements by the respective nucleotide (or IUPAC code, when polymorphic in the respective population), indels are marked by a dash. The translation frames refer to frame 1 that starts with ATG. Stop codons are marked by a star.

**Table 1 - supplement 1.**
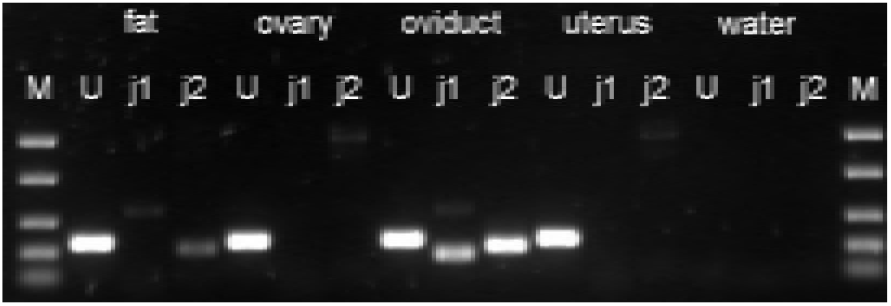
The ENCODE data do not provide the detail of expression in the different parts of the female reproductive system. Therefore, we have used RT-PCR across intron junctions to study *Udng3* expression in gonadal fat pad, ovary, oviduct, and uterus. Fat: gonadal fat pad; M: marker (from top to bottom: 1500 bp, 850 bp, 400 bp, 200 bp, 50 bp); U: *Uba1* (control gene, 255 bp); j1: *Udng3* junction 1 (161 bp); j2: *Udng3* junction 2 (209 bp).

